# Human Prostate Cancer-Associated Macrophage Subtypes with Prognostic Potential Revealed by Single-cell Transcriptomics

**DOI:** 10.1101/2020.06.19.160770

**Authors:** Joseph C Siefert, Bianca Cioni, Mauro J Muraro, Mohammed Alshalalfa, Judith Vivié, Henk van der Poel, Felix Y Feng, Lodewyk Wessels, Wilbert Zwart, Andries M Bergman

## Abstract

Macrophages in the tumor microenvironment are causally linked with prostate cancer development and progression, yet little is known about their composition in neoplastic human tissue. By performing single cell transcriptomic analysis of human prostate cancer resident macrophages, three distinct populations were identified in the diseased prostate. Unexpectedly, macrophages isolated from the tumor-adjacent site of the prostatectomy specimen were identical to those from the tumorous site. Markers associated with canonical M1 and M2 macrophage phenotypes were identifiable, however these were not the main factors defining unique subtypes. The genes selectively associated with each macrophage cluster were used to develop a gene signature which was highly associated with both recurrence-free and metastasis-free survival. These results highlight the relevance of tissue-specific macrophage subtypes in the tumour microenvironment for prostate cancer progression and demonstrates the utility of profiling single-cell transcriptomics in human tumor samples as a strategy to design gene classifiers for patient prognostication.

## Introduction

Blood-derived monocytes reach the majority of the tissues in the body, both cancer affected and normal, where they become tissue-resident macrophages [1]. These are extremely plastic and phenotypically heterogeneous immune cells, whose diversity is largely influenced by the microenvironment in which they reside [2]. Several studies showed that *in vitro* blood monocyte-derived macrophages can acquire a large spectrum of phenotypes depending on different stimuli present in the cell culture [3, 4]. However, these models do not capture the dynamic nature of macrophages in their native microenvironment. Human tissue-specific characterization of tumor associated macrophages (TAMs) is limited to glioma, skin and hepatocellular carcinoma [5-7], and there are no studies addressing prostate cancer (PCa)-specific macrophage phenotypic diversity.

The PCa tumor microenvironment (TME) is composed of various cells, including stromal, endothelial and immune cells, with tissue-resident macrophages representing one of the most predominant immune cell populations [8, 9]. Macrophages are critical mediators of tissue homeostasis and have the capacity to suppress cancer-associated processes, including tumor cell proliferation, angiogenesis and metastasis [10]. Multiple studies have shown a correlation between high infiltration of TAMs in the PCa microenvironment and poor prognosis, which suggests a role of these cells in cancer progression [11-15]. Given the prognostic significance of macrophages in the TME, strategies aiming to target these cells have emerged as strong candidates for cancer treatment [16-19].

It is also thought that immune cell type rather than sheer numbers of immune cells present in the TME relates to efficacy [20]. Various macrophage phenotypes have been described, including the pro-inflammatory/anti-tumor M1 state, and the anti-inflammatory/pro-tumor M2 state, both characterized by expression of specific markers [21]. However, macrophage diversity is likely not a binary division, but rather a continuum of phenotypes between M1 and M2 extremes [3, 4]. However, the diversity of human macrophage populations in PCa is not yet explored and, therefore, tissue specific markers of macrophage populations in prostate cancer are not yet defined.

To address this, we applied single-cell mRNA sequencing on myeloid cells isolated from prostatectomy specimens. Here we describe novel phenotypes of PCa associated macrophages and their distinct prognostic potential. Moreover, we propose new molecular markers for identification of these phenotypes in patients with localized disease. Understanding the unique macrophage populations in individual PCa and their effect on outcome will not only enhance our knowledge of prostate cancer biology and progression, but can better inform clinicians regarding a patients’ prognosis and treatment options.

## Materials and methods

### Patients, tumor specimens and Ethics Statement

Prostate biopsies were collected from post robotic-assisted laparoscopic prostatectomy (RALP) surgical specimen of PCa patients who did not receive any prior therapy (Gleason score 3+4 and 3+3). An average of three 18G biopsies were collected from both the tumorous and the adjacent non-tumorous site of the prostate, which were estimated by a pre-surgery multi parametric magnetic resonance imaging scan of the pelvis and palpation by the surgeon of the prostatectomy specimen. Fresh biopsies from the tumorous site and non-tumorous site of the prostatectomy specimen from 4 patients were processed separately for cell-surface markers CD14+ and/or CD11b+ myeloid cell isolation and submitted for single-cell RNA sequencing.

The use of patient data and biopsies from fresh prostatectomy specimens for research purposes at the Netherlands Cancer Institute have been executed pursuant to Dutch legislation and international standards. Prior to 25 May 2018, national legislation on data protection applied, as well as the International Guideline on Good Clinical Practice. From 25 May 2018 on, we also adhere to the GDPR. Within this framework, patients are informed and have always had the opportunity to object or actively consent to the (continued) use of their personal data & biospecimens in research. For the current study, informed consent was obtained from all patients. Hence, the procedures comply both with (inter-) national legislative and ethical standards.

### Tissue dissociation and CD11b+ and/or CD14+ cells sorting

Single-cell suspension was prepared from fresh PCa biopsies by mechanical dissociation within two hours after surgery. Biopsies were transported from the operation room on ice and minced with a scalpel in cold PBS + 0.5% BSA. Tissue was then mechanically dissociated using a gentleMACS™ Dissociator (MACS Milteny Biotec) using C-tubes (MACS Milteny Biotec) for 2 minutes as previously described [22]. Subsequently, the samples were filtered through a 70 μm strainer (BD Falcon) and spun down for 5 min at 300 g at 4°C. Cells were re-filtered through a 40 μm strainer (BD Falcon), spun down for 5 min at 300 g at 4°C and re-suspended in cold PBS + 0.5% BSA.

Cells of the dissociated biopsies were incubated with APC-CD45, PE-CD14, PE-CD11b and FITC-CD3 (all Ebioscience) for 20 minutes and washed before sorting using a Moflo Astrios (Beckman Coulter) or FACSAria IIu (BD BioSciences). As a first step, CD45+ leukocytes were selected, while small CD45+ cells (low SSC) were discarded as possible lymphocytes. Subsequently, CD14+ and/or CD11b+ single cells lacking CD3 expression were selected. Living single CD45+CD3-CD14+ and/or CD11b+ macrophages (based on DAPI and scatter properties) were sorted into eight 384 wells plates (Biorad) where cDNA synthesis was performed as previously described [23].

### Single-cell sequencing with SORT-seq

Single-cell mRNA sequencing was performed according to an adapted version of the SORT-seq protocol [23], using primers described by van den Brink *et al* [24]. In short, single cells were FACS sorted into 384-well plates containing 384 primers and Mineral oil (Sigma). After sorting, plates were snap-frozen on dry ice and stored at - 80°C. Subsequently, cells were heat-lysed at 65° C followed by cDNA synthesis using the CEL-Seq2 protocol [25] using robotic liquid handling platforms Nanodrop II (GC Biotech) and Mosquito (TTP Labtech). After second strand cDNA synthesis, the barcoded material was pooled into libraries of 384 cells and amplified using *in vitro* transcription [25]. Following amplification, the rest of the CEL-seq2 protocol was followed for preparation of the amplified cDNA library, using TruSeq small RNA primers (Illumina) as previously described [23]. The DNA library was paired-end sequenced on an Illumina Nextseq™500, high output, 1×75 bp.

### Single-cell sequencing data analysis

After Illumina sequencing, read 1 was assigned 26 base pairs and was used for identification of the Illumina library barcode and cell barcode. Unique molecular identifiers (UMI) tags were added to each read. These are molecular tags used in order to detect and quantify unique mRNA transcripts. More specifically, mRNA libraries were generated by fragmentation and reverse transcribed to cDNA with tag-specific primers. Read 2 was assigned 60 base pairs and used to map to the reference transcriptome of Hg19. Data was demultiplexed as described previously [26]. The Seurat v3.1.4 package was used in R v3.6.1 for processing the data [27]. Cells with less than 1000 features, greater than 6000 features, and greater than 1% mitochondrial reads were removed from analysis. The data were log-normalized with a scale factor of 10,000, and the 2000 most variable features were identified using a variance stabilizing transformation. The data was scaled according to all detected genes and principal component analysis was performed on the most variable genes. For linear dimensionality reduction, the number of principal components (20) was selected based on combined Jackstraw analysis, examination of elbow plots, difference in variation between subsequent principle components, and cumulative percent variation explained [28]. To identify clusters, a K-nearest neighbors graph was constructed from the selected principal components and clusters were identified from this using the Louvain algorithm at a resolution of 0.5 in Seurat. These were then projected with Uniform Manifold Approximation and Projection (UMAP) using uwot v0.1.5 and umap v0.2.4.1 packages in R. The reciprocal PCA method from Seurat v3 was used for data integration. In this procedure, the data from each patient was normalized, variable features were selected, the data was scaled, and principal component analysis was performed independently. The PCA space of each patient was then projected into each other patient to identify anchor points and integrate the datasets. The integrated data was then scaled and principal component analysis was performed as before.

### Differential expression and Gene Set Enrichment Analysis

Differential expression analysis was performed on normalized RNA values with minimum percentage (min.pct) and log fold-change (logfc) thresholds of 0.25 to identify only positive (upregulated) genes in each cluster. Significant differentially expressed genes were defined by Bonferroni adjusted p-value <0.05. Gene set enrichment analysis was performed using clusterProfiler v3.12.0 with msigdbr v7.1.1 database in R [29]. Hallmark gene set enrichment was performed by calculating logfc for all genes in each cluster as compared to the other two clusters, without any thresholds for min.pct or logfc, and ranking genes based on logfc.

### Generation and evaluation of gene signatures

Differentially expressed genes between macrophage clusters identified in the integrated dataset were used to construct prognostic signatures for biochemical relapse-free survival of PCa patients in a published MKSCC PCa dataset (GSE 21032) [30]. This dataset, comprised of 131 primary PCa patients with RNA expression and biochemical recurrence (BCR) as determined by serum prostate specific antigen (PSA) levels, was used to assess biochemical relapse-free survival (RFS). RFS was defined as time from prostatectomy to BCR (rise of PSA ≥0.2 ng/ml on two occasions). The gene signature (classifier) was generated employing Elastic net Cox regression using glmnet v4.0 in R [31]. The prognostic performance of the selected gene set was assessed in the MSKCC data by performing a nested 10-fold cross-validation (10FCV), with the full dataset split randomly split into 10 folds with each fold stratified for the number of events and Gleason score. The Elastic net regularization parameters (alpha and lambda) were optimized as follows. A preselected set of values for alpha were tested (0 to 1 by 0.1). For each value of alpha in this set, we performed 10FCV to determine the optimal value of lambda. To this end, we selected the value of lambda associated with the minimum, average mean-squared error (MSE) across the folds. This procedure delivered, for each value of alpha, the optimal value of lambda and the associated MSE. We then selected the value of alpha that delivered the lowest MSE across all values of alpha in the set. Hazard ratios (HR), confidence intervals (CI), p-values, and Harrel’s C-index (concordance index) for the MSKCC evaluation were generated using the coxph function in the survival v3.1-12 package in R [32]. Survival plots were made by selecting the high (top 25%), low (bottom 25%) and intermediate (middle 50%) risk of relapse from the CV predictions, fit with event censoring and BCR-free time from MSKCC dataset using the survfit function in the survival package. Receiver operating characteristic (ROC) curves and AUC were calculated using predictions from the CV with ROCR v1.0-7 in R [33]. The final macrophage gene signature used for validation was generated by fitting the full MSKCC dataset with the optimized parameter values.

### Validation of gene signatures in independent cohorts

Gene signatures were tested in three independent cohorts by first extracting the coefficients (betas) for the selected genes from the model fit, then multiplying the scaled gene expression data in the independent datasets by these coefficients. The prospective Decipher cohort contains anonymized genome-wide expression profiles from clinical use of the Decipher test in the radical prostatectomy (RP) setting, between February 2014 to August 2017, retrieved from the Decipher GRID™ (NCT02609269). The retrospective natural history cohort from Johns Hopkins Medical Institute is comprised of men treated with RP, with a median follow up time of 108 months [34]. The Mayo clinic cohort is a retrospective cohort of men treated with RP, with a median follow-up time of 156 months [35, 36]. Model discriminatory capability was assessed based on the AUC. Cox proportional hazards modeling was used to estimate the Hazard Ratio of metastasis-free survival for patients with high signature (top25%).

### Public availability of data

Limited and specific single cell RNA sequencing data of patient macrophages can be found at GSE133094. RNA expression data for Mayo (GSE46691, GSE62116) and JHMI (GSE79957) cohorts are also available.

## Results

### Single-cell analysis of myeloid cells isolated from PCa biopsies

To identify the macrophage populations present in diseased human prostates, fresh biopsies were collected from ‘tumorous’ and ‘tumor-adjacent’ areas of post radical prostatectomy (RP) specimens. Four previously untreated PCa patients, aged 50-74 years, with an initial serum PSA between 7.6 and 9.3 ng/ml and diagnosed with a Gleason score 6-7 and a clinical stage pT2-3 adenocarcinoma of the prostate were included in this study (Supplementary Table 1). The procedure for obtaining native PCa associated myeloid cells is depicted in Figure 1A. Tissue resident macrophages were isolated from the biopsies by successively FACS sorting a single-cell suspension of the biopsies for CD45+ leukocytes, followed by isolation of CD3-CD14+ and/or CD11b+ myeloid cells [37-39]. In total, 1920 cells, including 911 cells isolated from the tumorous and 1009 cells isolated from the tumor-adjacent areas of the prostates were sequenced on 8 plates (Supplementary Table 1). Cells with less than 2000 UMIs were not considered, while only genes that were detected with at least 4 UMIs in at least 3 cells, were used for further analysis. In plates 4 and 7, very few cells above the 2000 UMI cut-off were found. Furthermore, high levels of technical artefact genes like KCNQ1OT1, which is a non-coding region rich in poly-A repeats and often found in cell transcripts of poor quality were also detected [40]. For these reasons, plates 4 and 7 were excluded from further analysis. After additional quality control filtering to remove low-quality cells and doublets/multiplets, plate 8 was found to contain very few cells and was also removed from analysis. The distributions of the remaining cells were normal and 751 cells were retained for subsequent analysis (Supplementary Figure 1).

**Figure 1.**
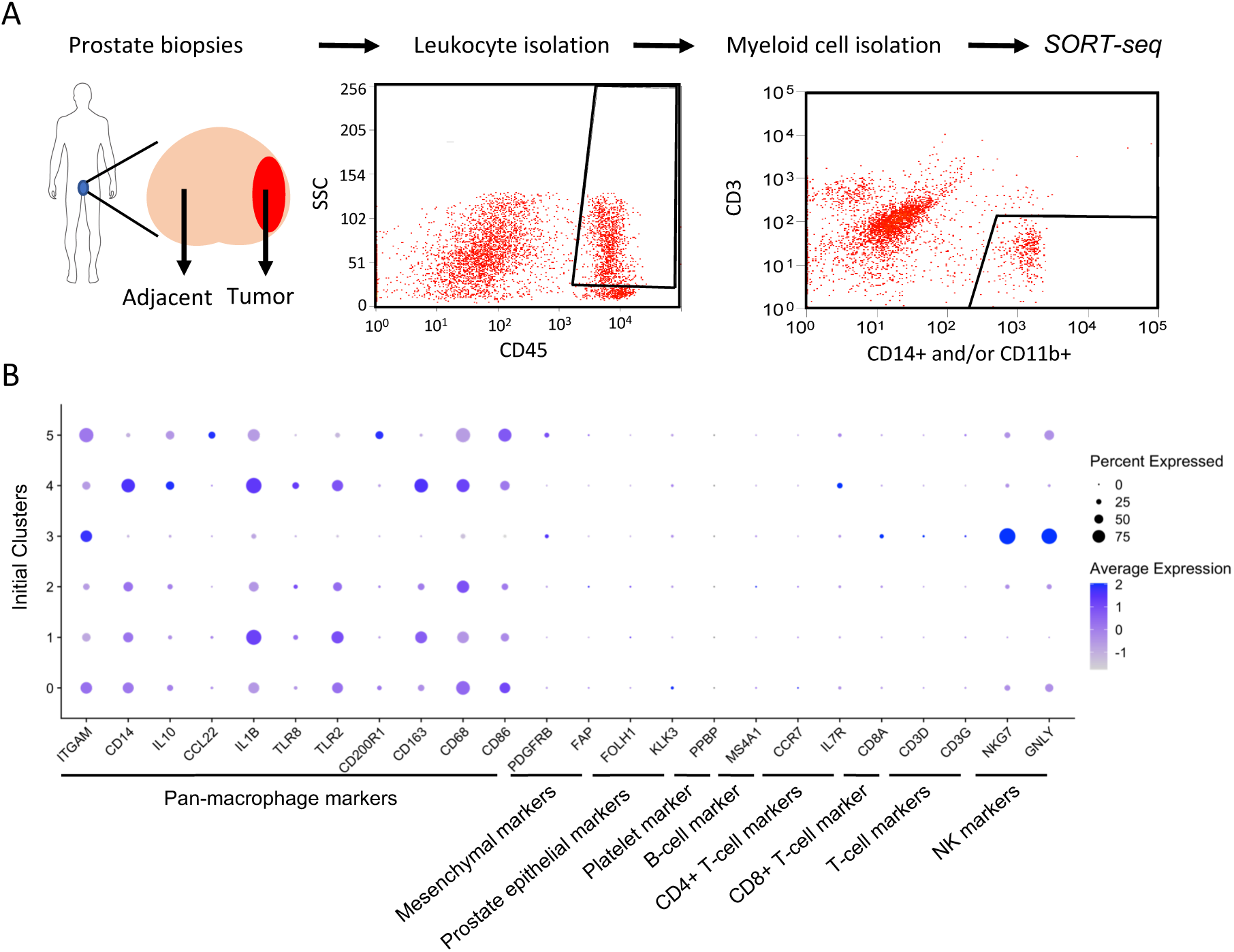
Single cell sequencing of myeloid cells from prostate cancer patients. (A) Experimental procedure: single cell suspensions of multiple biopsies from the tumor and tumor-adjacent portions of radical prostatectomy specimens were sequentially FACS sorted to isolate CD45+/CD14+ and/or CD45+/CD11b+ myeloid cells, after negative selection for CD45+CD3+ lymphocytes. Subsequently, single cells were processed for SORT-seq analysis. (B) Dot plot showing markers for macrophages, NK cells (NKG7 & GNLY), and other immune and prostate cell types present in the initial six clusters identified. Cells were positively identified as macrophages and NK cells.

### Clustering of myeloid cells to identify PCa macrophage subtypes

To surmount the implicit noise of individual features in scRNA data, principal component (PC) analysis was used to reduce dimensionality, followed by graph-based clustering to identify populations of highly-interconnected cells [27]. Initial clustering of the data yielded 6 independent clusters, with cluster 3 being substantially divergent from the remaining clusters (Supplementary Figure 2A). This was also evident in the first principal component (Supplementary Figure 2B). Examination of the genes within PC1, as well as markers for various cell types across all clusters revealed that cells in cluster 3 expressed the well-known natural killer (NK) cell markers NKG7 and GNLY (Figure 1B, Supplementary Figures 2C-D) [41]. The presence of these cells after FACS sorting is likely residual from the CD11b (ITGAM) selection (Figure 1B). Since the focus of this study is macrophages, these NK cells were removed from further analysis. The 641 cells in the remaining clusters are considered macrophages as they all express various pan-macrophage markers (CD68, CD86), while lacking expression of established markers for other immune cell types (T-cell, B-cell, NK-cell), prostate epithelium (FOLH1, KLK3), and mesenchymal cells (PDGFRB, FAP) (Figure 1B) [42, 43].

**Figure 2.**
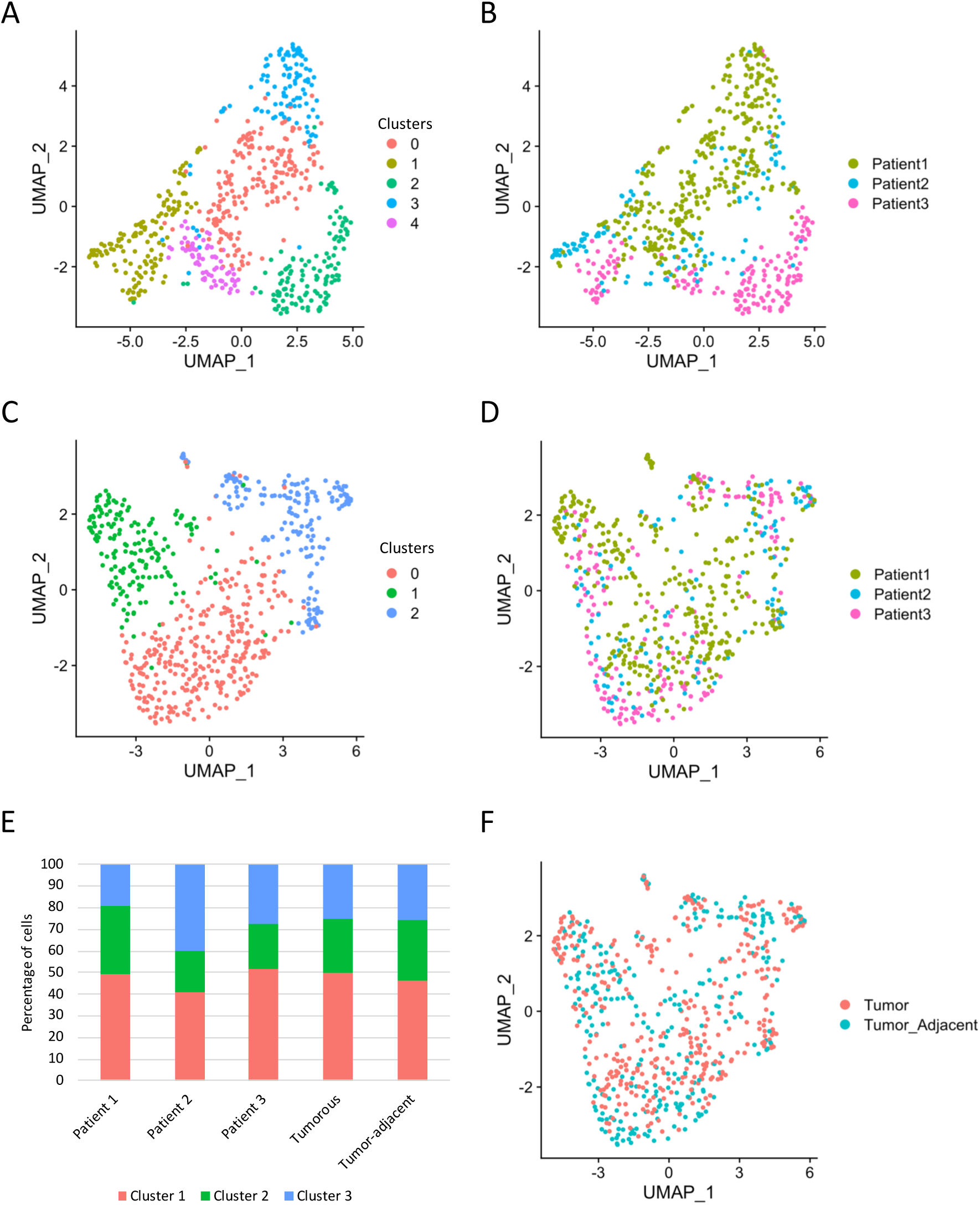
Identification of prostate cancer-associated macrophage subtypes. (A) UMAP projections of macrophage clustering after NK cell removal. Colors indicate five distinct clusters. (B) UMAP projections of cells from each patient reveals strong patient-specific batch effects. Each color represents cells from a different patient. (C) UMAP projections after reciprocal-PCA integration removing batch effects between patients. Colors indicate three distinct clusters. (D) UMAP projection of integrated data shows all patient samples are comparable and do not cluster separately. Each color represents cells from a different patient. (E) Distributions of cells within each cluster, with percentage of cells from each cluster show on the y-axis. (F) UMAP projection of macrophages isolated from tumor (pink) and tumor adjacent (green) biopsies are comparable and do not cluster separately.

After removal of NK cells, the remaining macrophages were reanalyzed as above, and 20 PCs were selected for further analysis (Supplementary Figure 3). Clustering these PCs yielded 5 populations, however the clustering was highly patient specific (Figures 2A-B). To remove these patient-specific batch effects, a reciprocal PCA method was employed to integrate the patient datasets [27]. Clustering of the final integrated dataset revealed 3 distinct macrophage subtypes (Figure 2C). The cells from each patient were no longer forming isolated or dominant clusters, but were instead distributed across all three clusters, indicating that the reciprocal PCA method was effective at removing the patient-specific batch effects (Figures 2D-E). Unexpectedly, the macrophages from the tumor and tumor-adjacent biopsies showed identical distributions between the clusters, suggesting that there are no differences between the macrophages present in the tumorous and tumor-adjacent prostate tissue (Figures 2E-F). Cumulatively these results indicate that there are three biologically distinct macrophage subpopulations present throughout the diseased human prostate.

**Figure 3.**
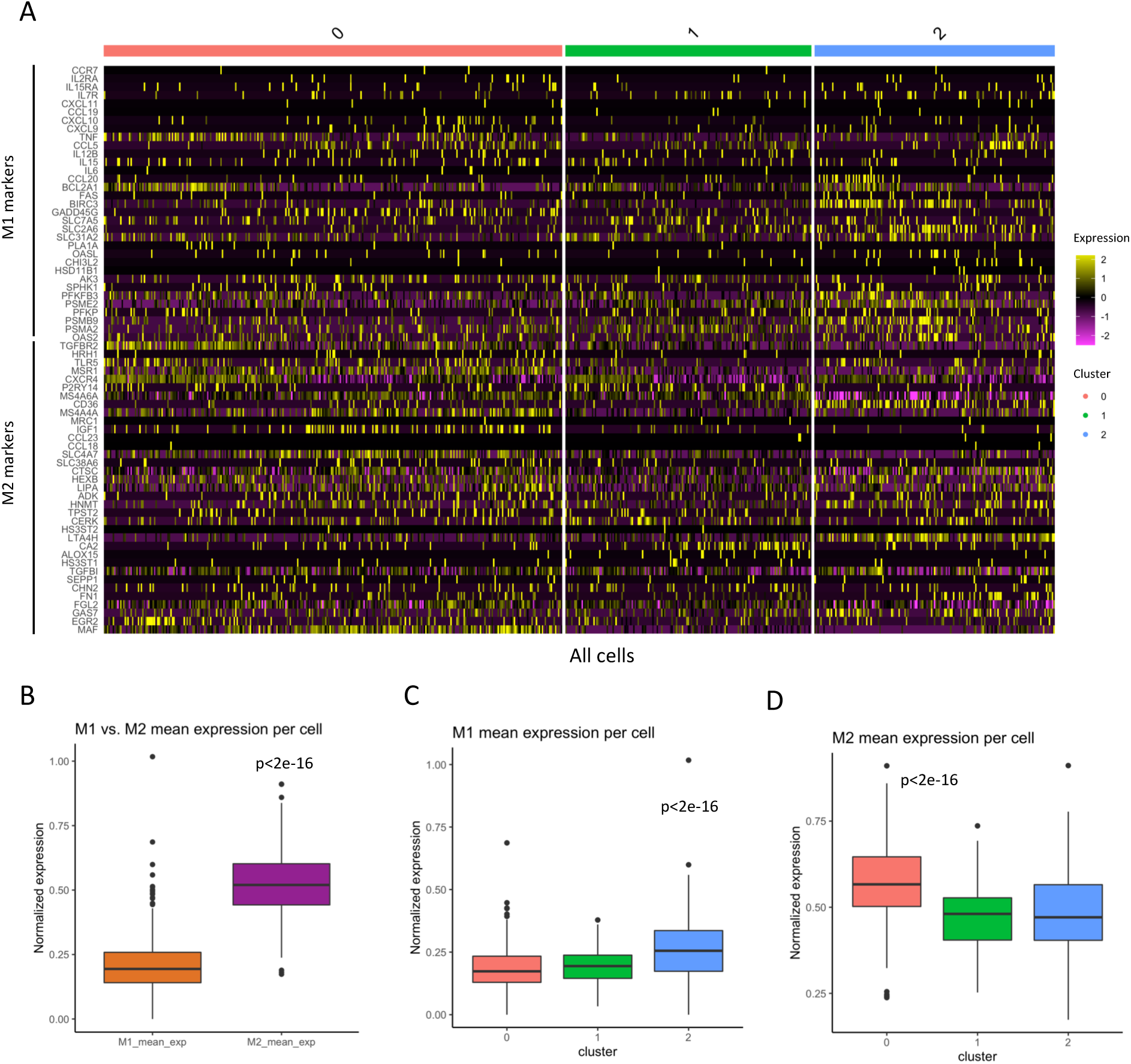
Canonical M1 and M2 macrophage marker expression in prostate cancer associated macrophages. (A) Heatmap of all detectable M1 and M2 macrophage markers in relation to the three macrophage clusters. Color bar indicates RNA expression level scaled by z-score. (B) Mean expression of all M1 (orange) or M2 (purple) markers from (A) averaged together within each cell. ***M1 vs M2; p<2e-16 Welch’s t-test. (C) Average normalized expression of all M1 markers in (A) per cell within each cluster. ***p<2e-16 One-way ANOVA with post-hoc Tukey HSD. (D) Average normalized expression of all M2 markers in (A) per cell within each cluster; ***p<2e-16 One-way ANOVA with post-hoc Tukey HSD.

### Evaluation of M1 and M2 macrophage phenotypes in PCa clusters

As a first step to examine the identity of these macrophage clusters, previously established markers associated with M1-like and M2-like phenotypes were investigated [21]. Plotting all detectable M1 and M2 marker genes for each cell in a heatmap revealed that many markers are not readily detectable in all cells and there is no clear M1/M2 separation between these clusters (Figure 3A). Given the varying expression levels and the sparsity of marker expression, averaging individual markers within each cluster was not useful in evaluating M1/M2 identity within the clusters, though it appeared that M2 markers were generally expressed at higher levels (Supplementary Figures 4A-B). For these reasons, all M1 and M2 markers were separately combined by averaging the RNA expression of all M1 or M2 markers within each cell. From this analysis it is evident that the mean expression level of all combined M2 markers per cell are higher than the mean M1 marker expression levels (Figure 3B, Supplementary Figure 4C). Furthermore, the mean M1 expression levels were slightly but significantly higher in cluster 2, while the mean M2 expression levels were significantly elevated in cluster 0 (Figures 3C-D). These results demonstrate that while a slightly elevated expression of the averaged M1 and M2 markers can be detected in certain clusters, these are not the main factors contributing to the variation that separates these macrophage populations.

**Figure 4.**
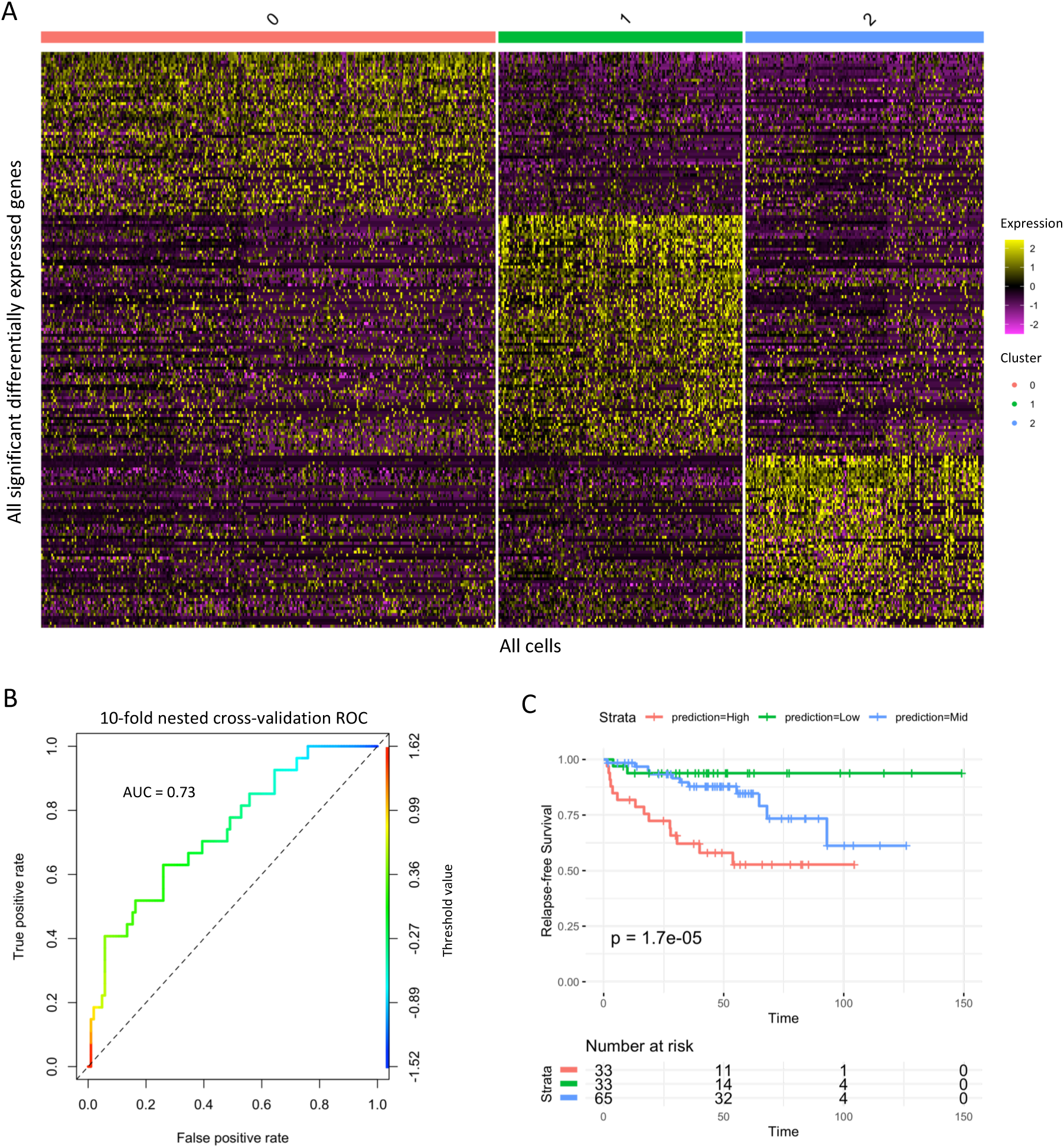
Gene signature from differentially expressed macrophage genes has significant prognostic value. (A) Heatmap of all significant differentially expressed genes present in the integrated dataset by macrophage cluster (Bonferroni adjusted p-value <0.05). Color bar indicates integrated expression level scaled by z-score. (B) Macrophage gene signature outcome predictions from 10-fold cross-validation in MSKCC dataset. Colors indicate patient subpopulations of predicted high (red; 25%), medium (blue; 50%) and low (green;25%) risk of recurrence. p-value is from log-rank test. (C) Receiver operating characteristic (ROC) curve for 10-fold cross-validation of macrophage signature in MSKCC dataset, colorized by threshold value.

### Identification of differentially expressed genes and biological pathways

To determine the biological differences between these three macrophage clusters, differential expression analysis was performed to detect marker genes in each cluster. This analysis identified 468 significantly differentially expressed genes, with 164 genes identified as markers in cluster 0, 199 genes in cluster 1, and 105 genes in cluster 2 (Figure 4A, Supplementary Table 2). The most differentially expressed genes from each cluster show either expression only in their cluster, or elevated expression as compared to the other clusters (Supplementary Figure 5A-B). Examining this list of genes showed only 11/68 M1 and M2 markers to be differentially expressed, with 3/33 M1 markers upregulated in cluster 2 and 6/35 M2 markers upregulated in cluster 0 (Supplementary Table 3). These results agree with the slight enrichments observed in Figures 3C-D, however there are also two M2 markers upregulated in cluster 2, further exemplifying the need for better stratification.

**Figure 5.**
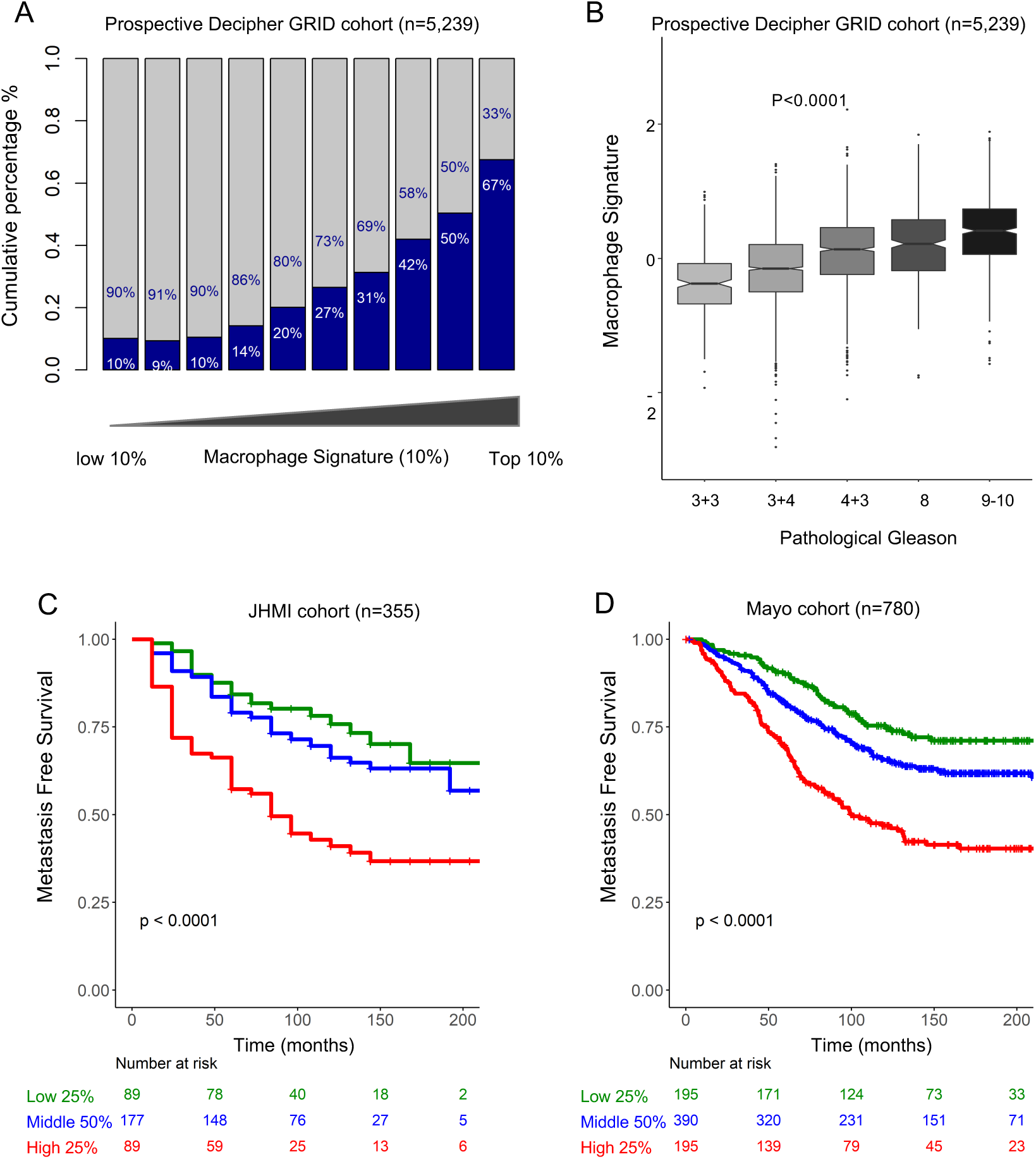
Validation of the gene signature in independent Prostate cancer cohorts. (A) Macrophage gene signature predictions correlate in a prospective cohort of >5000 patients with high Decipher GRID scores (blue) vs. low and intermediate scores (grey). (B) Macrophage gene signature predictions correlate with pathological Gleason score in Decipher GRID. ****p-value <2e-16 from linear model between Gleason and Macrophage signature. (C-D) Gene signature outcome predictions in two independent PCa cohorts from JHMI (C) and Mayo Clinic (D), stratifying patients on low (green, bottom 25%), medium (blue; middle 50%) or high (red; top 25%) risk of metastatic disease. Y-axis shows metastasis-free survival, number of patients at risk are indicated, p-values from log-rank test.

To explore the functional pathways that genes associated with each cluster are involved in, gene set enrichment analysis (GSEA) was performed (Supplementary Figure 6). Cluster 0 genes showed activation of the hallmark TNFα signalling via NFKB as well as WNT β-catenin signalling, and suppression of interferon pathways (IFN-α and IFN-γ), MTORC1 signalling, and complement pathways, among others (Supplemental Figure 6A). Conversely, cluster 2 showed activation of multiple inflammatory pathways including IFN-α, IFN-γ, TNFα, and complement, while showing suppression of WNT β-catenin signalling and cell cycle pathways (Supplemental Figure 6B). Cluster 1, however, showed suppression of multiple immune pathways (IFN-α, IFN-γ, TNFα, among others), and activation of cell cycle pathways (E2F targets, MYC targets, G2M checkpoint) as well as MTORC1 signalling (Supplementary Figure 6C). To explore the possibility of PCa TAM regulation of T-cells, known markers of T-cell regulation by TAMs were interrogated in the data [44]. Very few of these markers were readily detectable and only one, CSF1R, was found to be significantly differentially expressed (Supplementary Figure 7). Collectively, these results suggest that each macrophage population is involved in unique biological functions.

### Generation and evaluation of macrophage gene signature

Using the differentially expressed genes in the integrated dataset, a prospective gene signature was developed using a published PCa dataset from Memorial Sloan Kettering Cancer Center (MSKCC) [27]. The performance of the macrophage gene signature was evaluated employing 10-fold nested cross-validation on the MSKCC dataset by receiver-operator characteristic (ROC) as performance measure (Figure 4B). As expected, in a Cox regression analysis this classifier showed a significant association with relapse-free survival (hazard ratio (HR) = 4.1, p = 1.7e-05) (Figure 4C). Furthermore, in a multi-variate analysis including the signature with Gleason score (biopsy and pathological), pre-diagnosis biopsy PSA levels, seminal vesicle invasion (SVI), extracapsular extension (ECE), and clinical stage the signature was found to be an independent predictor of outcome (Table 1).

### Validation of gene signature in independent PCa cohorts

To further assess the prognostic value of the macrophage gene signature, it was validated in three independent cohorts from the Decipher GRID registry. The first cohort is a prospective Decipher GRID cohort containing RNA expression data from >5,000 radical prostatectomy (RP) patients and includes basic demographic and pathological data, but not longitudinal clinical outcomes. This cohort was used to associate the signature to Decipher risk groups and pathological Gleason score (Figure 5A-B). Since this cohort has no metastasis outcome yet, high Decipher group was used as a surrogate of metastasis potential since it was heavily validated for that endpoint [34, 36, 45]. The second cohort is a retrospective natural history cohort (n=355) comprised of men treated with RP at Johns Hopkins Medical Institutions (JHMI) [34]. The third cohort is a retrospective cohort (n=780) of men treated with RP at the Mayo Clinic [36, 45]. All three cohorts are described in Supplementary Table 4.

The strength of association of the macrophage signature with metastasis-free survival was tested using a Cox regression analysis on the Mayo and JHMI cohorts and the signature showed significant association with metastasis-free survival (Mayo: HR = 1.89, p-value = 1.0e-06; JHMI: HR = 2.25, p-value = 3.3e-05) (Figure 5C-D). In both cohorts, the classifier was also found to be an independent predictor of metastasis in multivariate analysis (Table 2). Taken together, these results indicate that profiling single-cell RNA expression in PCa associated macrophages and identifying subpopulations present in the diseased prostate can have significant prognostic value in predicting patients’ likelihood of biochemical relapse and metastasis. These results lay the foundation for profiling macrophage populations in prostate cancer and other cancer types, and will inform future studies investigating the immune systems’ role in cancer progression.

## Discussion

Macrophages can either promote or suppress cancer development and progression depending on their specific phenotype and function. In this study, we defined the degree of human PCa-specific macrophage diversity through single-cell sequencing with the aim to identify PCa-specific macrophage populations. Three macrophage subtypes were identified, and while some canonical M1 and M2 markers were present, these were not adequate to define the clusters. Using the genes differentially expressed in each cluster we were able to develop a gene signature with significant prognostic value in multiple PCa cohorts. Remarkably, no difference was observed between macrophages collected from the tumor and the adjacent non-tumor sites, suggesting that tumorigenic factors may also affect distant non-tumorigenic sites. Furthermore, this suggests that these macrophage populations could in theory be detected from a biopsy regardless to tumor cell percentage. This is important because the prognostic value of the gene signature outweighs the biopsy Gleason score and pre-diagnosis biopsy PSA levels, and is approaching the significance of pathological Gleason, suggesting a possible path to identifying high-risk patients without necessitating radical prostatectomy. Additionally, the macrophage signature and pathological Gleason score were both independent predictors in our multivariate analysis, suggesting that the signature can provide additional prognostic information. However, these finding will require further experimental validation before such measures could be employed.

The gene set enrichment analysis performed in this study suggests that each macrophage subtypes is involved in unique biological processes. Cluster 1 does not appear to be participating in inflammatory pathways and may represent a proliferative feeder cell type, or otherwise less differentiated macrophage subtype. Cluster 0 appears to be largely anti-inflammatory, while cluster 2 appears primarily pro-inflammatory. These results agree with the notion that macrophages can broadly adopt either a pro-inflammatory or anti-inflammatory phenotype, and this could either potentiate or mediate cancer progression [3, 4, 21]. It will be important for future studies to explore the role these macrophage populations play in prostate cancer, and to investigate targeting these subtypes and their associated pathways with immunotherapy.

Cancer immunotherapies, specifically those inhibiting T cell immune checkpoints, have generated significant impact in recent years, with established efficacy in advanced melanoma [46, 47], non-small cell lung cancer [48, 49] and bladder cancer patients [50, 51]. However, in other cancers, including prostate cancer (PCa), immunotherapy efficacy is limited [52, 53]. The uncertain therapeutic efficacy of immunotherapy in PCa is partly due to a poor infiltration of immune cells in the TME [13, 54-57]. Moreover, TAMs display an ability to modulate tumor immunity by suppression of T cell recruitment and function, though the precise mechanisms have yet to be elucidated [44]. Several direct and indirect suppressive actions of macrophages on T cell functions have been suggested, including involvement of immune checkpoints ligands (*e*.*g*.: PDL1, B7-H4), cytokines (*e*.*g*.: IL-10, CXCL10, CCL22) and cell surface receptors (*e*.*g*.: CD206, CSF1R) [44]. However, in this study, only the colony-stimulating factor 1 receptor (CSF1R), which is a key regulator of immunosuppressive macrophage expansion, was found to be enriched in cluster 0. Whether the macrophage subtypes discovered in this study play a role in T cell regulation will be an important question for future studies.

Limitations of this study include the small number of patients included in the study and the absence of assessment of protein expression of the key selected genes. To this end, future studies should include immunohistochemistry analysis to further support our findings.

In conclusion, in this study we demonstrate the relevance of using single-cell transcriptomics from PCa-associated macrophages as a prognostication strategy for individual patients. We propose that targeting unique tumor-associated macrophage subtypes, as opposed to all macrophages, can provide a therapeutic avenue to combat prostate cancer and potentially other cancer types.

## Acknowledgements

We thank our collaborators from Single Cell Discoveries B.V. supported by the Hubrecht Institute and the Oncode Institute.

## Funding details

This work was supported by the FP7 MCA-ITN Grant agreement 317445 – TIMCC

## Disclosure of Potential Conflicts of Interest

The authors report no conflict of interest

## Notes

### Competing Interest Statement

The authors have declared no competing interest.

## References

1. Davies, L.C., et al., Tissue-resident macrophages. Nat Immunol, 2013. 14(10): p. 986–95.

2. Gordon, S., A. Pluddemann, and F. Martinez Estrada, Macrophage heterogeneity in tissues: phenotypic diversity and functions. Immunol Rev, 2014. 262(1): p. 36–55.

3. Beyer, M., et al., High-resolution transcriptome of human macrophages. PLoS One, 2012. 7(9): p. e45466.

4. Xue, J., et al., Transcriptome-based network analysis reveals a spectrum model of human macrophage activation. Immunity, 2014. 40(2): p. 274–88.

5. Muller, S., et al., Single-cell profiling of human gliomas reveals macrophage ontogeny as a basis for regional differences in macrophage activation in the tumor microenvironment. Genome Biol, 2017. 18(1): p. 234.

6. Dan Xue, T.T., Christina Morse and Robert Lafyatis, Single-cell RNA sequencing reveals different subsets of macrophage and dendritic cells in human skin. J Immunol 2019. 202(1).

7. Zhang, Q., et al., Landscape and Dynamics of Single Immune Cells in Hepatocellular Carcinoma. Cell, 2019. 179(4): p. 829–845 e20.

8. Poltavets, V., et al., The Role of the Extracellular Matrix and Its Molecular and Cellular Regulators in Cancer Cell Plasticity. Front Oncol, 2018. 8: p. 431.

9. Mantovani, A., et al., Tumour-associated macrophages as treatment targets in oncology. Nat Rev Clin Oncol, 2017. 14(7): p. 399–416.

10. Poh, A.R. and M. Ernst, Targeting Macrophages in Cancer: From Bench to Bedside. Front Oncol, 2018. 8: p. 49.

11. Lanciotti, M., et al., The role of M1 and M2 macrophages in prostate cancer in relation to extracapsular tumor extension and biochemical recurrence after radical prostatectomy. Biomed Res Int, 2014. 2014: p. 486798.

12. Shimura, S., et al., Reduced infiltration of tumor-associated macrophages in human prostate cancer: association with cancer progression. Cancer Res, 2000. 60(20): p. 5857–61.

13. Nonomura, N., et al., Infiltration of tumour-associated macrophages in prostate biopsy specimens is predictive of disease progression after hormonal therapy for prostate cancer. BJU Int, 2011. 107(12): p. 1918–22.

14. Hu, W., et al., Alternatively activated macrophages are associated with metastasis and poor prognosis in prostate adenocarcinoma. Oncol Lett, 2015. 10(3): p. 1390–1396.

15. Gollapudi, K., et al., Association between tumor-associated macrophage infiltration, high grade prostate cancer, and biochemical recurrence after radical prostatectomy. Am J Cancer Res, 2013. 3(5): p. 523–9.

16. Escamilla, J., et al., CSF1 receptor targeting in prostate cancer reverses macrophage-mediated resistance to androgen blockade therapy. Cancer Res, 2015. 75(6): p. 950–62.

17. Xu, J., et al., CSF1R signaling blockade stanches tumor-infiltrating myeloid cells and improves the efficacy of radiotherapy in prostate cancer. Cancer Res, 2013. 73(9): p. 2782–94.

18. Wong, D., et al., Targeting CXCR4 with CTCE-9908 inhibits prostate tumor metastasis. BMC Urol, 2014. 14: p. 12.

19. Huang, E.H., et al., A CXCR4 antagonist CTCE-9908 inhibits primary tumor growth and metastasis of breast cancer. J Surg Res, 2009. 155(2): p. 231–6.

20. Higano, C.S., et al., Integrated data from 2 randomized, double-blind, placebo-controlled, phase 3 trials of active cellular immunotherapy with sipuleucel-T in advanced prostate cancer. Cancer, 2009. 115(16): p. 3670–9.

21. Martinez, F.O. and S. Gordon, The M1 and M2 paradigm of macrophage activation: time for reassessment. F1000Prime Rep, 2014. 6: p. 13.

22. Norstrom, M.M., et al., Novel method to characterize immune cells from human prostate tissue. Prostate, 2014. 74(14): p. 1391–9.

23. Muraro, M.J., et al., A Single-Cell Transcriptome Atlas of the Human Pancreas. Cell Syst, 2016. 3(4): p. 385–394 e3.

24. van den Brink, S.C., et al., Single-cell sequencing reveals dissociation-induced gene expression in tissue subpopulations. Nat Methods, 2017. 14(10): p. 935–936.

25. Hashimshony, T., et al., CEL-Seq2: sensitive highly-multiplexed single-cell RNA-Seq. Genome Biol, 2016. 17: p. 77.

26. Grun, D., L. Kester, and A. van Oudenaarden, Validation of noise models for single-cell transcriptomics. Nat Methods, 2014. 11(6): p. 637–40.

27. Stuart, T., et al., Comprehensive Integration of Single-Cell Data. Cell, 2019. 177(7): p. 1888–1902 e21.

28. Chung, N.C. and J.D. Storey, Statistical significance of variables driving systematic variation in high-dimensional data. Bioinformatics, 2015. 31(4): p. 545–54.

29. Yu, G., et al., clusterProfiler: an R package for comparing biological themes among gene clusters. OMICS, 2012. 16(5): p. 284–7.

30. Taylor, B.S., et al., Integrative genomic profiling of human prostate cancer. Cancer Cell, 2010. 18(1): p. 11–22.

31. Friedman, J., T. Hastie, and R. Tibshirani, Regularization Paths for Generalized Linear Models via Coordinate Descent. J Stat Softw, 2010. 33(1): p. 1–22.

32. Terry M. Therneau, P.M.G., Modeling Survival Data: Extending the Cox Model, in Springer. 2000, Springer: New York.

33. Sing, T., et al., ROCR: visualizing classifier performance in R. Bioinformatics, 2005. 21(20): p. 3940–1.

34. Ross, A.E., et al., Tissue-based Genomics Augments Post-prostatectomy Risk Stratification in a Natural History Cohort of Intermediate- and High-Risk Men. Eur Urol, 2016. 69(1): p. 157–65.

35. Erho, N., et al., Discovery and validation of a prostate cancer genomic classifier that predicts early metastasis following radical prostatectomy. PLoS One, 2013. 8(6): p. e66855.

36. Karnes, R.J., et al., Validation of a genomic classifier that predicts metastasis following radical prostatectomy in an at risk patient population. J Urol, 2013. 190(6): p. 2047–53.

37. Perera, P.Y., et al., CD11b/CD18 acts in concert with CD14 and Toll-like receptor (TLR) 4 to elicit full lipopolysaccharide and taxol-inducible gene expression. J Immunol, 2001. 166(1): p. 574–81.

38. Lasitschka, F., et al., Human monocytes downregulate innate response receptors following exposure to the microbial metabolite n-butyrate. Immun Inflamm Dis, 2017. 5(4): p. 480–492.

39. da Silva, T.A., et al., CD14 is critical for TLR2-mediated M1 macrophage activation triggered by N-glycan recognition. Sci Rep, 2017. 7(1): p. 7083.

40. Zhu, S., X.O. Zhang, and L. Yang, Panning for Long Noncoding RNAs. Biomolecules, 2013. 3(1): p. 226–41.

41. Butler, A., et al., Integrating single-cell transcriptomic data across different conditions, technologies, and species. Nat Biotechnol, 2018. 36(5): p. 411–420.

42. Sugimoto, H., et al., Identification of fibroblast heterogeneity in the tumor microenvironment. Cancer Biol Ther, 2006. 5(12): p. 1640–6.

43. Fearon, D.T., The carcinoma-associated fibroblast expressing fibroblast activation protein and escape from immune surveillance. Cancer Immunol Res, 2014. 2(3): p. 187–93.

44. DeNardo, D.G. and B. Ruffell, Macrophages as regulators of tumour immunity and immunotherapy. Nat Rev Immunol, 2019. 19(6): p. 369–382.

45. Howard LE et al., Validation of a genomic classifier for prediction of metastasis and prostate cancer-specific mortality in African-American men following radical prostatectomy in an equal access healthcare setting. Prostate Cancer Prostatic Dis., 2019.

46. Hodi, F.S., et al., Improved survival with ipilimumab in patients with metastatic melanoma. N Engl J Med, 2010. 363(8): p. 711–23.

47. Robert, C., et al., Ipilimumab plus dacarbazine for previously untreated metastatic melanoma. N Engl J Med, 2011. 364(26): p. 2517–26.

48. Reck, M., et al., Pembrolizumab versus Chemotherapy for PD-L1-Positive Non-Small-Cell Lung Cancer. N Engl J Med, 2016. 375(19): p. 1823–1833.

49. Antonia, S.J., et al., Durvalumab after Chemoradiotherapy in Stage III Non-Small-Cell Lung Cancer. N Engl J Med, 2017. 377(20): p. 1919–1929.

50. Ardolino, L. and A. Joshua, Immune checkpoint inhibitors in malignancy. Aust Prescr, 2019. 42(2): p. 62–67.

51. Bellmunt, J., et al., Pembrolizumab as Second-Line Therapy for Advanced Urothelial Carcinoma. N Engl J Med, 2017. 376(11): p. 1015–1026.

52. Kwon, E.D., et al., Ipilimumab versus placebo after radiotherapy in patients with metastatic castration-resistant prostate cancer that had progressed after docetaxel chemotherapy (CA184-043): a multicentre, randomised, double-blind, phase 3 trial. Lancet Oncol, 2014. 15(7): p. 700–12.

53. Beer, T.M., et al., Randomized, Double-Blind, Phase III Trial of Ipilimumab Versus Placebo in Asymptomatic or Minimally Symptomatic Patients With Metastatic Chemotherapy-Naive Castration-Resistant Prostate Cancer. J Clin Oncol, 2017. 35(1): p. 40–47.

54. Vitkin, N., et al., The Tumor Immune Contexture of Prostate Cancer. Front Immunol, 2019. 10: p. 603.

55. Kiniwa, Y., et al., CD8+ Foxp3+ regulatory T cells mediate immunosuppression in prostate cancer. Clin Cancer Res, 2007. 13(23): p. 6947–58.

56. Strasner, A. and M. Karin, Immune Infiltration and Prostate Cancer. Front Oncol, 2015. 5: p. 128.

57. Ebelt, K., et al., Prostate cancer lesions are surrounded by FOXP3+, PD-1+ and B7-H1+ lymphocyte clusters. Eur J Cancer, 2009. 45(9): p. 1664–72.

