## Supplementary Information for "Human Prostate Cancer-Associated Macrophage Subtypes with Prognostic Potential Revealed by Single-cell Transcriptomics"

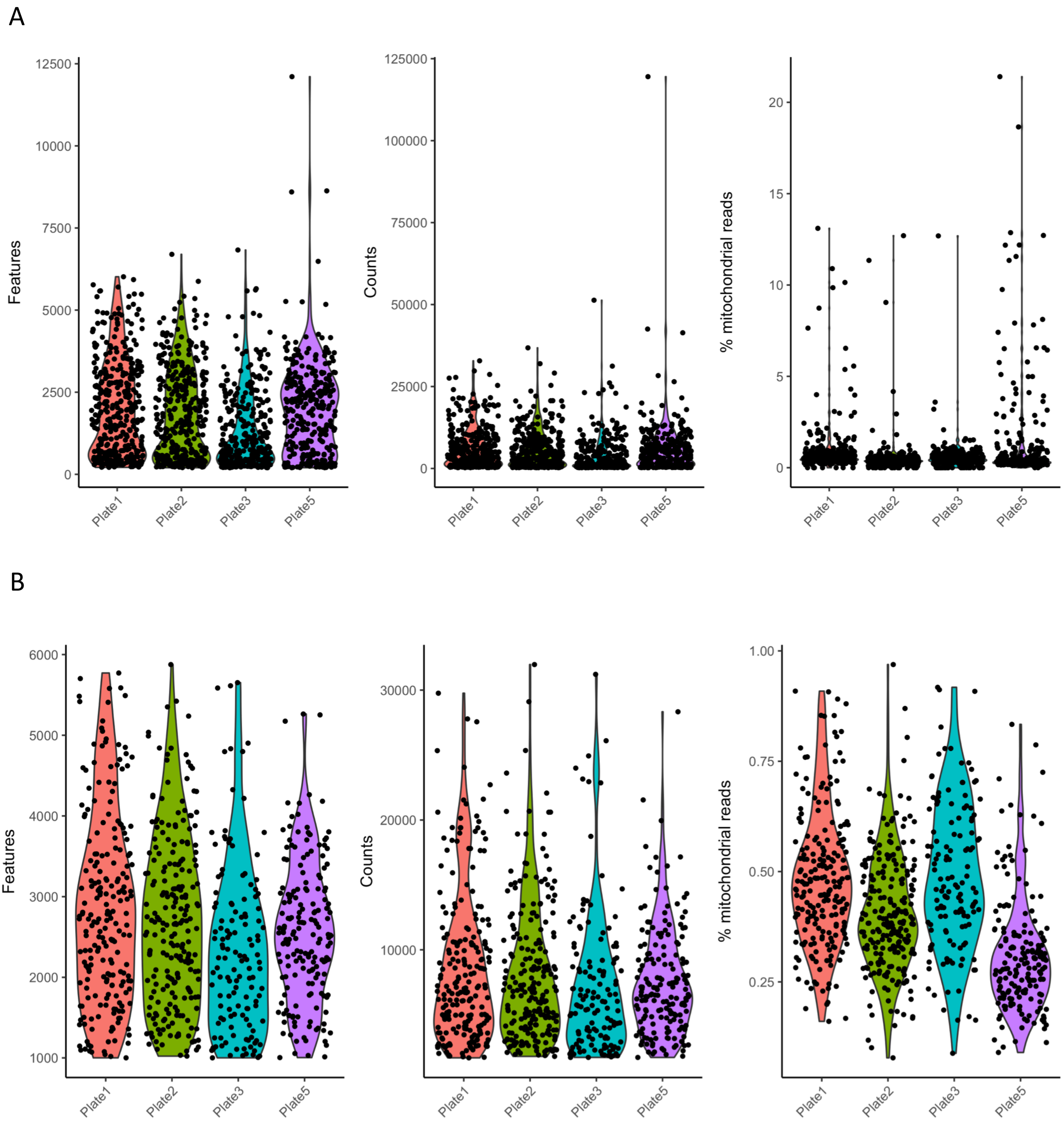


**Supplementary Figure 1. Quality control metrics of single-cell RNA-seq data.** (A) Plot showing the number of features (genes), RNA counts, and percentage mitochondrial reads of the data set prior to filtering. (B) Quality control metrics as in (A) after filtering.

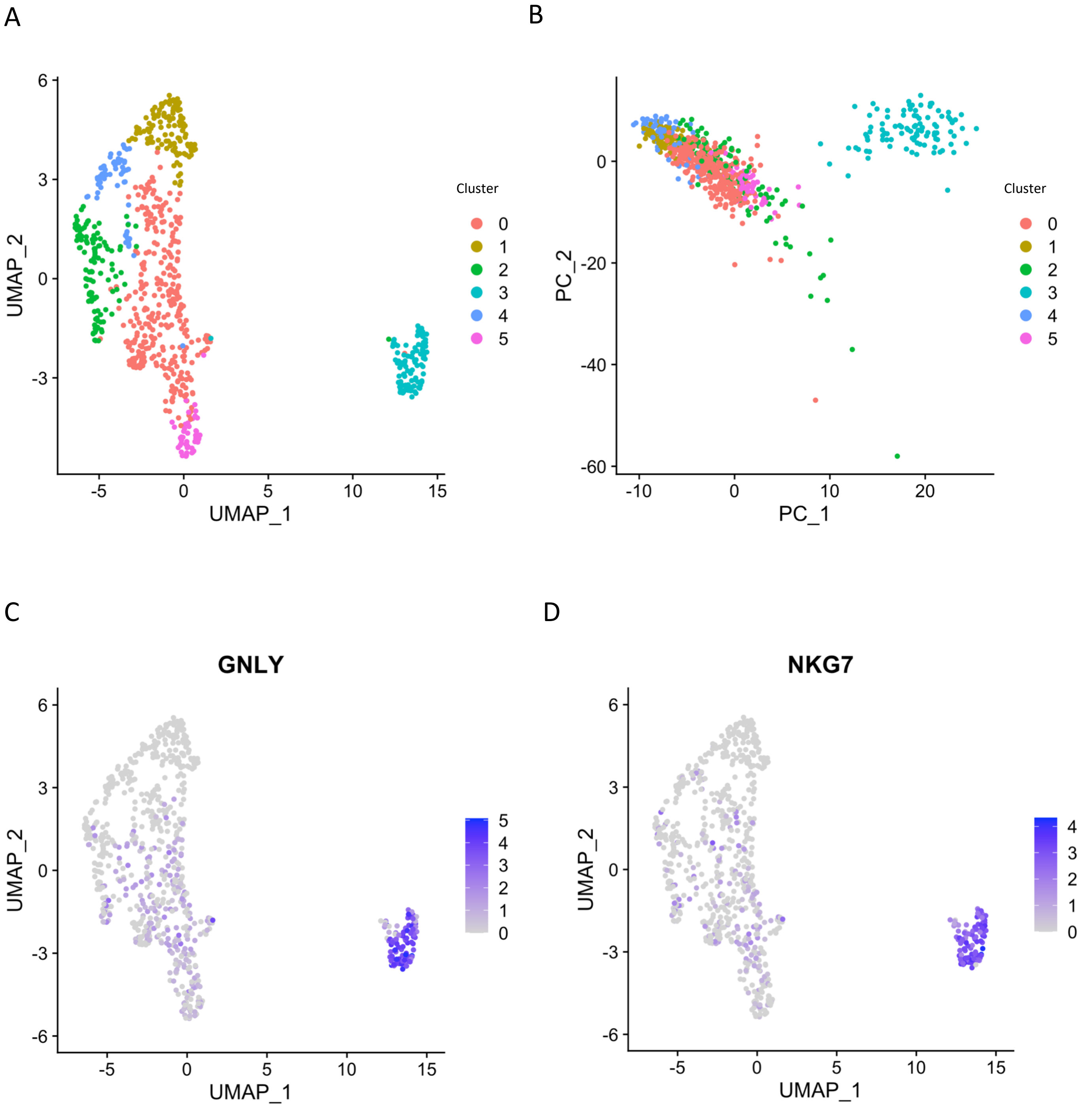


**Supplementary Figure 2. Identification of contaminating cells.** (A) UMAP projection of filtered dataset colored by the six identified clusters. (B) First two principal components (PC) showing the difference in cluster 3 for PC1. (C-D) UMAP projections showing high expression of the natural killer (NK) cell markers GNLY (C) and NKG7 (D) in cluster 3. Color bar indicates normalized expression values.

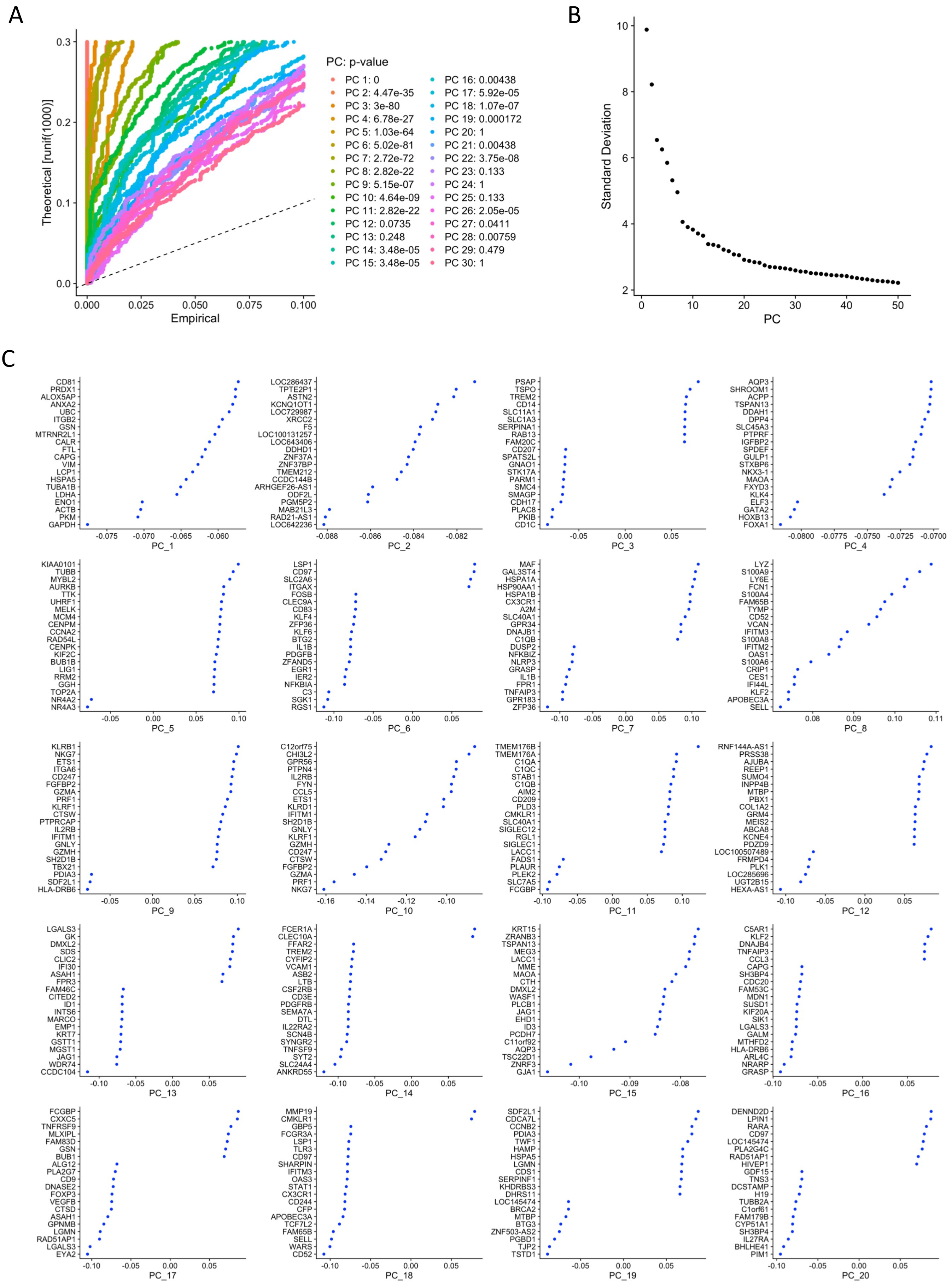


**Supplementary Figure 3. Principal component selection.** (A) Jackstraw plot showing analysis of the first 30 principal components (PC), with each PC indicated by a different color. (B) Elbow plot showing the standard deviation for each of the 50 analyzed PCs. (C) Top 20 genes contributing to the variation observed in each of the 20 selected PCs.

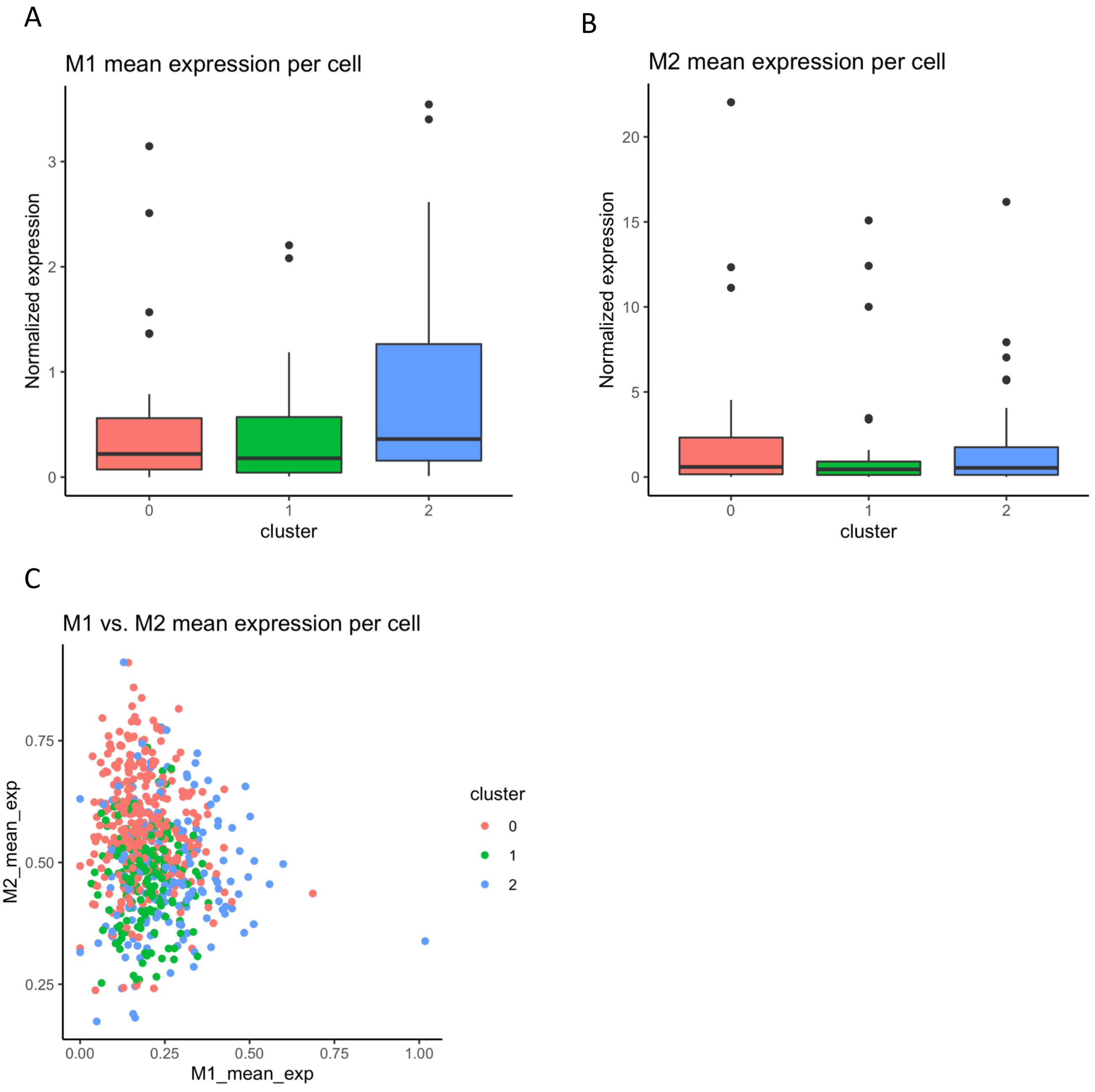


**Supplementary Figure 4. Canonical M1 and M2 macrophage markers expression throughout the clusters.** (A) Mean expression of each individual M1 marker per cluster. (B) Mean expression of each individual M2 marker per cluster. (C) Averaged expression of all M1 and M2 markers in each cell colored by cluster, plotting mean expression of M1 markers (x-axis) over mean expression of M2 markers (y-axis).

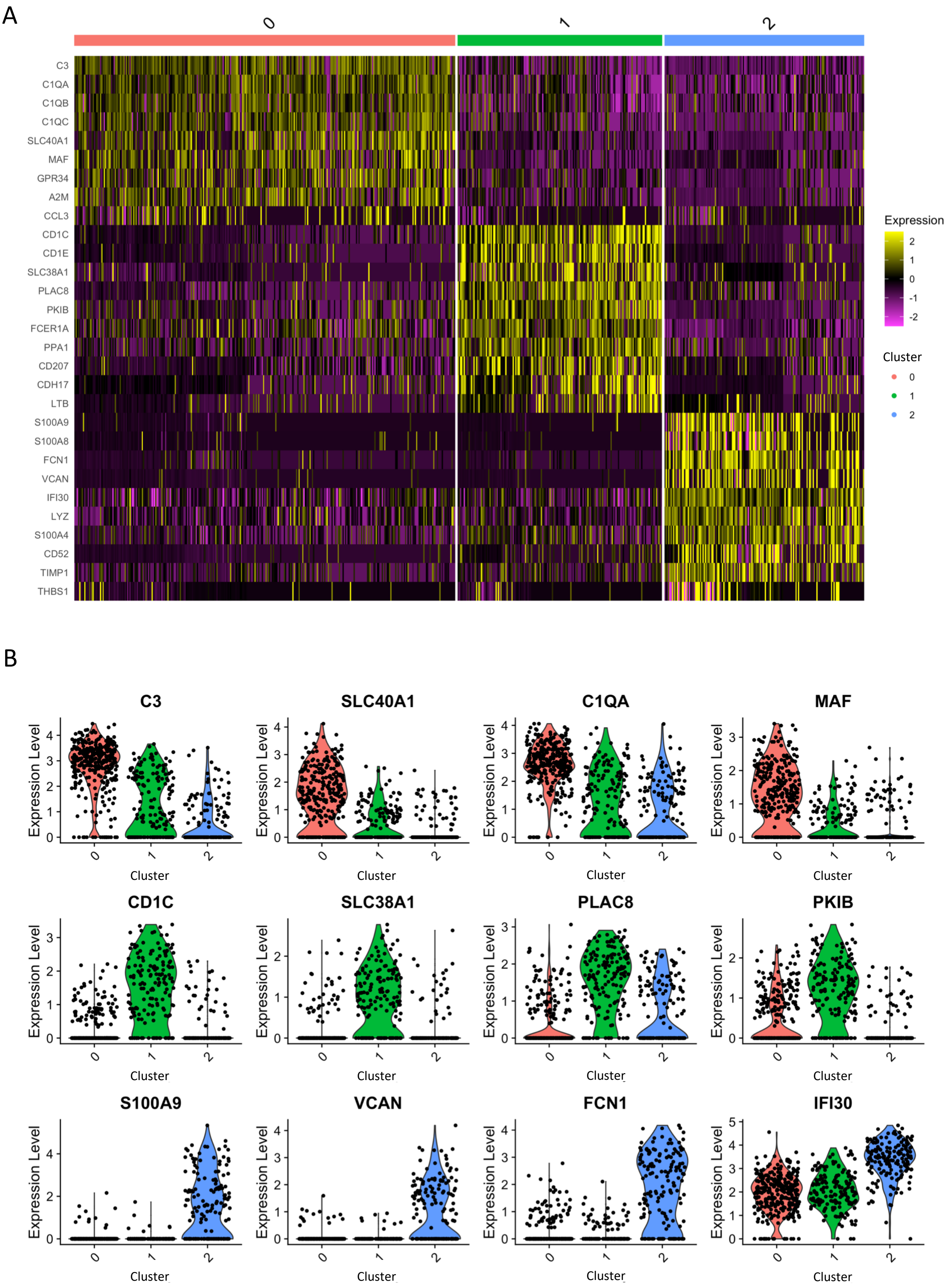


**Supplementary Figure 5. Most differentially expressed genes in each cluster.** (A) Heatmap of the top 10 most differentially expressed genes in each macrophage cluster with expression level scaled to z-score. (B) Violin plots for representative differentially expressed genes in each cluster showing normalized RNA expression levels. Colors indicate macrophage clusters and each black dot represents a cell.

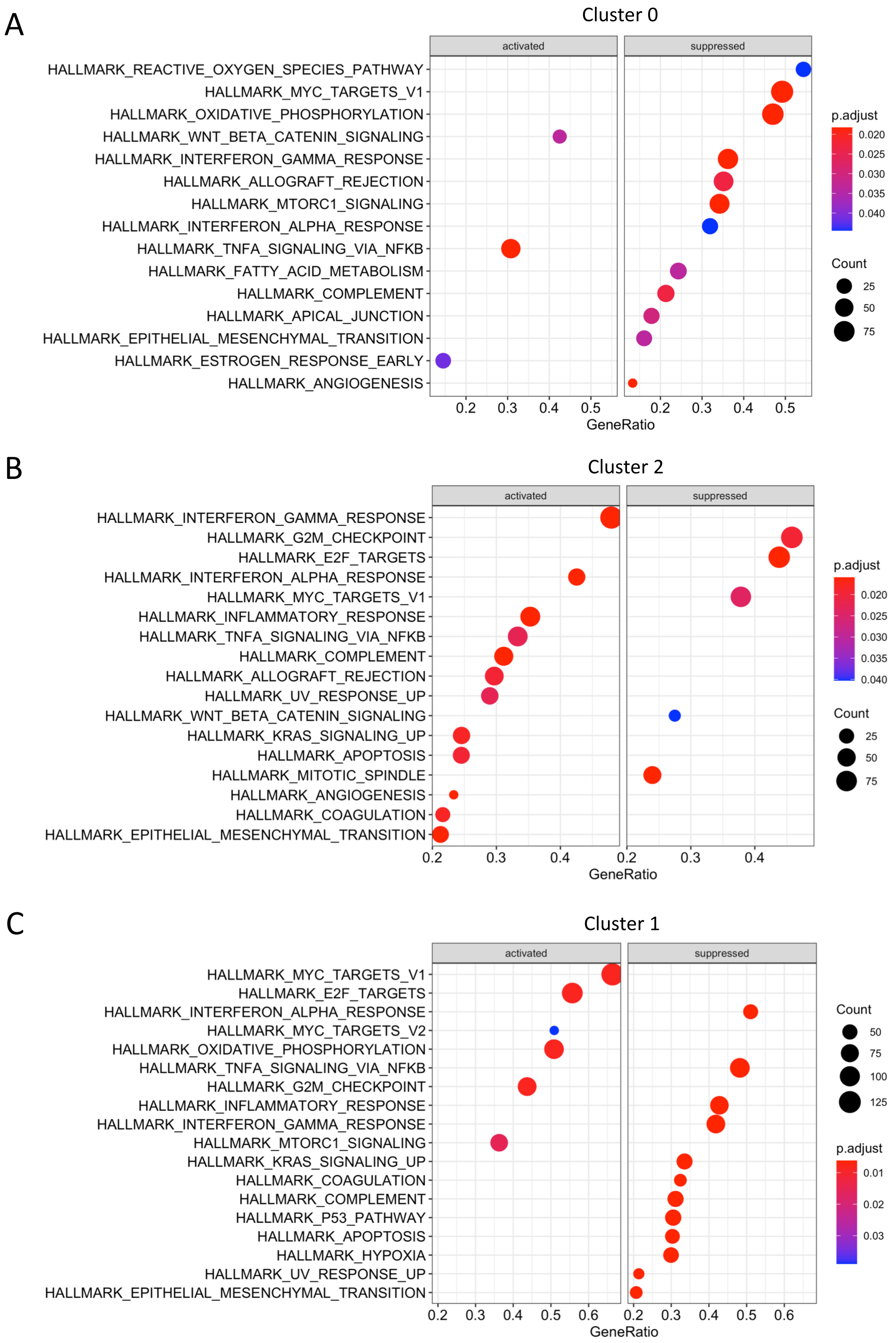


**Supplementary Figure 6. Gene set enrichment analysis for each cluster.** (A) Gene set enrichment analysis (GSEA) for hallmark gene sets in cluster 0. (B) GSEA for hallmark gene sets in cluster 2. (C) GSEA for hallmark gene sets in cluster 1. Circle size indicates number of genes identified in the gene set, color shows Benjamini-Hochberg adjusted p-value.

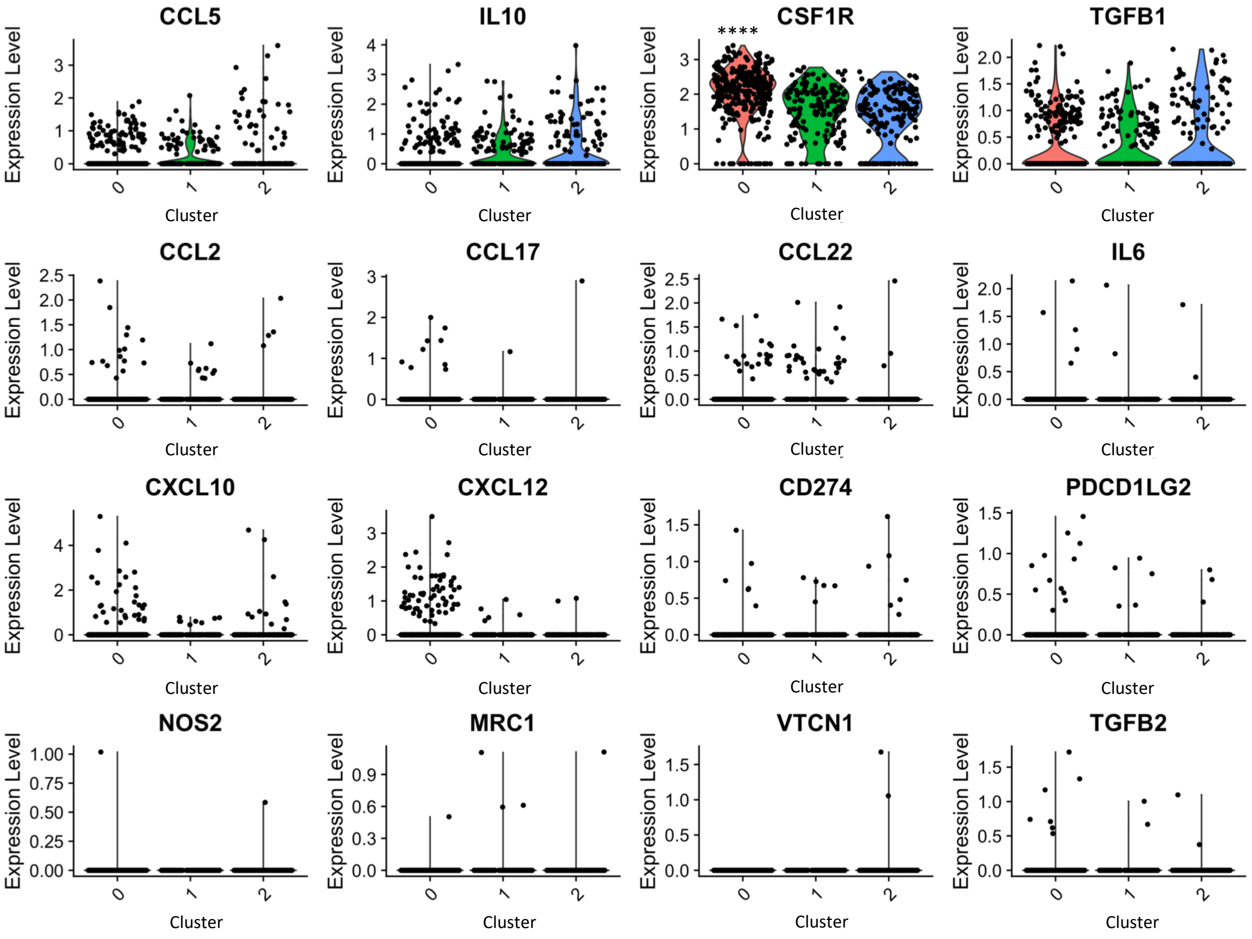


**Supplementary Figure 7. Expression of markers for TAM regulation of T-cells.** RNA expression levels for selected markers of tumor-associated macrophage regulation of T-cells. ****Bonferroni adjusted p-value <1e-16.

**Supplementary Table 2.** All significant differentially expressed genes between clusters

| gene | p_val | avg_logFC | pct.1 | pct.2 | p_val_adj | cluster |
|---|---|---|---|---|---|---|
| C3 | 2.43E-64 | 1.531 | 0.942 | 0.518 | 4.22E-60 | 0 |
| SLCO2B1 | 2.03E-55 | 1.213 | 0.891 | 0.415 | 3.53E-51 | 0 |
| C1QA | 4.95E-55 | 1.118 | 0.968 | 0.591 | 8.59E-51 | 0 |
| ITM2B | 1.64E-50 | 0.738 | 0.997 | 0.976 | 2.84E-46 | 0 |
| C1QB | 1.14E-46 | 1.077 | 0.955 | 0.642 | 1.99E-42 | 0 |
| C1QC | 1.71E-45 | 1.049 | 0.894 | 0.448 | 2.96E-41 | 0 |
| SLC40A1 | 3.42E-39 | 1.35 | 0.756 | 0.348 | 5.94E-35 | 0 |
| MAF | 3.06E-37 | 1.083 | 0.756 | 0.327 | 5.32E-33 | 0 |
| GPR34 | 8.77E-37 | 0.942 | 0.768 | 0.379 | 1.52E-32 | 0 |
| A2M | 8.63E-36 | 1.038 | 0.797 | 0.403 | 1.50E-31 | 0 |
| MARCKS | 3.33E-34 | 0.777 | 0.939 | 0.782 | 5.77E-30 | 0 |
| PLXDC1 | 2.64E-32 | 0.863 | 0.662 | 0.258 | 4.58E-28 | 0 |
| RGS1 | 1.29E-29 | 0.903 | 0.997 | 0.873 | 2.24E-25 | 0 |
| PDGFB | 1.72E-29 | 0.916 | 0.682 | 0.291 | 2.99E-25 | 0 |
| CSF1R | 8.56E-29 | 0.599 | 0.945 | 0.842 | 1.49E-24 | 0 |
| IGSF21 | 1.04E-27 | 0.643 | 0.431 | 0.07 | 1.80E-23 | 0 |
| LPAR6 | 3.23E-27 | 0.741 | 0.839 | 0.567 | 5.61E-23 | 0 |
| SORL1 | 5.58E-27 | 0.785 | 0.868 | 0.664 | 9.69E-23 | 0 |
| USP53 | 7.99E-26 | 0.852 | 0.695 | 0.367 | 1.39E-21 | 0 |
| GGTA1P | 8.39E-26 | 0.733 | 0.666 | 0.345 | 1.46E-21 | 0 |
| BIN1 | 1.08E-25 | 0.688 | 0.691 | 0.345 | 1.88E-21 | 0 |
| FCGR3A | 5.74E-25 | 0.667 | 0.849 | 0.521 | 9.97E-21 | 0 |
| MS4A7 | 1.01E-24 | 0.642 | 0.91 | 0.761 | 1.75E-20 | 0 |
| TREM2 | 1.55E-24 | 0.628 | 0.518 | 0.139 | 2.69E-20 | 0 |
| CD74 | 1.64E-24 | 0.324 | 1 | 1 | 2.84E-20 | 0 |
| EPB41L2 | 2.56E-24 | 0.692 | 0.801 | 0.539 | 4.45E-20 | 0 |
| MEF2A | 2.92E-24 | 0.616 | 0.868 | 0.624 | 5.07E-20 | 0 |
| FRMD4A | 4.17E-23 | 0.652 | 0.595 | 0.273 | 7.23E-19 | 0 |
| BLNK | 3.99E-22 | 0.558 | 0.441 | 0.121 | 6.92E-18 | 0 |
| HERPUD1 | 7.62E-22 | 0.54 | 0.99 | 0.936 | 1.32E-17 | 0 |
| CEBPD | 1.27E-21 | 0.622 | 0.955 | 0.827 | 2.21E-17 | 0 |
| CLEC9A | 1.43E-21 | 0.615 | 0.431 | 0.106 | 2.48E-17 | 0 |
| KCTD12 | 2.38E-21 | 0.491 | 0.981 | 0.915 | 4.13E-17 | 0 |
| DAB2 | 3.39E-21 | 0.693 | 0.72 | 0.448 | 5.88E-17 | 0 |
| ZFHX3 | 5.32E-21 | 0.642 | 0.768 | 0.533 | 9.23E-17 | 0 |
| CX3CR1 | 1.49E-20 | 0.74 | 0.862 | 0.67 | 2.59E-16 | 0 |
| DOCK4 | 1.86E-20 | 0.701 | 0.675 | 0.409 | 3.22E-16 | 0 |
| BHLHE41 | 4.43E-19 | 0.631 | 0.624 | 0.309 | 7.68E-15 | 0 |
| SGK1 | 5.96E-19 | 0.916 | 0.907 | 0.803 | 1.03E-14 | 0 |
| CD81 | 7.11E-19 | 0.392 | 0.99 | 0.833 | 1.23E-14 | 0 |
| SESN1 | 8.43E-19 | 0.53 | 0.698 | 0.421 | 1.46E-14 | 0 |

| | | | | | | |
|---|---|---|---|---|---|---|
| HLA-DPA1 | 1.31E-18 | 0.316 | 1 | 1 | 2.27E-14 | 0 |
| RAPGEF6 | 1.62E-16 | 0.447 | 0.559 | 0.291 | 2.81E-12 | 0 |
| IFNGR1 | 2.90E-16 | 0.422 | 0.949 | 0.839 | 5.03E-12 | 0 |
| AXL | 3.54E-16 | 0.472 | 0.852 | 0.585 | 6.15E-12 | 0 |
| SRGAP1 | 6.42E-16 | 0.612 | 0.553 | 0.264 | 1.11E-11 | 0 |
| OLFML3 | 8.16E-16 | 0.576 | 0.415 | 0.145 | 1.42E-11 | 0 |
| LTC4S | 8.83E-16 | 0.461 | 0.408 | 0.145 | 1.53E-11 | 0 |
| ST6GAL1 | 8.87E-16 | 0.565 | 0.675 | 0.461 | 1.54E-11 | 0 |
| C3AR1 | 1.05E-15 | 0.461 | 0.743 | 0.521 | 1.83E-11 | 0 |
| MSR1 | 1.06E-15 | 0.496 | 0.685 | 0.436 | 1.83E-11 | 0 |
| JUN | 5.02E-15 | 0.594 | 0.932 | 0.803 | 8.71E-11 | 0 |
| ADAM28 | 1.40E-14 | 0.447 | 0.759 | 0.503 | 2.43E-10 | 0 |
| TGFBR2 | 4.52E-14 | 0.53 | 0.698 | 0.5 | 7.85E-10 | 0 |
| FCGR2A | 4.57E-14 | 0.493 | 0.723 | 0.482 | 7.93E-10 | 0 |
| UNC5B | 4.94E-14 | 0.475 | 0.492 | 0.245 | 8.58E-10 | 0 |
| SFMBT2 | 6.66E-14 | 0.517 | 0.624 | 0.391 | 1.16E-09 | 0 |
| GAL3ST4 | 7.24E-14 | 0.509 | 0.45 | 0.209 | 1.26E-09 | 0 |
| RB1 | 1.12E-13 | 0.459 | 0.669 | 0.452 | 1.95E-09 | 0 |
| MEF2C | 1.12E-13 | 0.455 | 0.768 | 0.552 | 1.95E-09 | 0 |
| GPR155 | 2.39E-13 | 0.515 | 0.672 | 0.467 | 4.16E-09 | 0 |
| SLC4A7 | 3.03E-13 | 0.493 | 0.669 | 0.455 | 5.26E-09 | 0 |
| PELI1 | 3.64E-13 | 0.483 | 0.598 | 0.37 | 6.31E-09 | 0 |
| RNASET2 | 7.35E-13 | 0.332 | 0.955 | 0.894 | 1.28E-08 | 0 |
| HLA-DMA | 8.53E-13 | 0.325 | 0.958 | 0.852 | 1.48E-08 | 0 |
| CLEC7A | 8.57E-13 | 0.402 | 0.871 | 0.782 | 1.49E-08 | 0 |
| MGAT4A | 1.01E-12 | 0.406 | 0.749 | 0.561 | 1.75E-08 | 0 |
| MCF2L | 1.59E-12 | 0.354 | 0.277 | 0.07 | 2.77E-08 | 0 |
| MERTK | 1.68E-12 | 0.518 | 0.444 | 0.209 | 2.92E-08 | 0 |
| HLA-DRB1 | 2.12E-12 | 0.279 | 1 | 0.997 | 3.68E-08 | 0 |
| NEAT1 | 2.91E-12 | 0.6 | 0.807 | 0.667 | 5.04E-08 | 0 |
| ZNF618 | 3.41E-12 | 0.515 | 0.547 | 0.321 | 5.93E-08 | 0 |
| ZFP36L2 | 4.00E-12 | 0.424 | 0.994 | 0.921 | 6.95E-08 | 0 |
| ADORA3 | 4.72E-12 | 0.466 | 0.569 | 0.339 | 8.19E-08 | 0 |
| AKR1B1 | 1.06E-11 | 0.406 | 0.72 | 0.509 | 1.85E-07 | 0 |
| GLUL | 1.30E-11 | 0.31 | 0.929 | 0.842 | 2.25E-07 | 0 |
| HLA-DPB1 | 1.65E-11 | 0.266 | 1 | 0.982 | 2.87E-07 | 0 |
| PLXDC2 | 2.20E-11 | 0.44 | 0.572 | 0.364 | 3.82E-07 | 0 |
| WASF2 | 2.83E-11 | 0.409 | 0.859 | 0.767 | 4.90E-07 | 0 |
| VSIG4 | 3.84E-11 | 0.436 | 0.592 | 0.348 | 6.67E-07 | 0 |
| MS4A4A | 4.67E-11 | 0.471 | 0.614 | 0.412 | 8.10E-07 | 0 |
| MAN2A1 | 5.86E-11 | 0.342 | 0.65 | 0.433 | 1.02E-06 | 0 |
| SPRED1 | 6.90E-11 | 0.45 | 0.534 | 0.327 | 1.20E-06 | 0 |
| WLS | 8.07E-11 | 0.31 | 0.251 | 0.067 | 1.40E-06 | 0 |

| | | | | | | |
|---|---|---|---|---|---|---|
| ARHGAP24 | 9.00E-11 | 0.336 | 0.283 | 0.091 | 1.56E-06 | 0 |
| NFKBIA | 1.04E-10 | 0.277 | 0.987 | 0.903 | 1.80E-06 | 0 |
| GPX1 | 1.08E-10 | 0.298 | 0.984 | 0.936 | 1.87E-06 | 0 |
| SLC1A3 | 1.53E-10 | 0.501 | 0.495 | 0.279 | 2.66E-06 | 0 |
| OGFRL1 | 2.51E-10 | 0.388 | 0.707 | 0.515 | 4.35E-06 | 0 |
| CXCL16 | 2.70E-10 | 0.358 | 0.868 | 0.748 | 4.69E-06 | 0 |
| ABCA1 | 2.72E-10 | 0.411 | 0.36 | 0.155 | 4.72E-06 | 0 |
| VASH1 | 2.85E-10 | 0.419 | 0.72 | 0.558 | 4.95E-06 | 0 |
| TM6SF1 | 2.86E-10 | 0.382 | 0.598 | 0.421 | 4.97E-06 | 0 |
| FMNL3 | 2.90E-10 | 0.404 | 0.595 | 0.391 | 5.03E-06 | 0 |
| TIAM1 | 3.40E-10 | 0.388 | 0.444 | 0.23 | 5.89E-06 | 0 |
| IGF1 | 4.57E-10 | 0.451 | 0.264 | 0.088 | 7.94E-06 | 0 |
| PTGS1 | 9.21E-10 | 0.409 | 0.662 | 0.509 | 1.60E-05 | 0 |
| QKI | 9.30E-10 | 0.351 | 0.868 | 0.8 | 1.61E-05 | 0 |
| MAST3 | 9.71E-10 | 0.391 | 0.415 | 0.212 | 1.69E-05 | 0 |
| CREG1 | 1.18E-09 | 0.36 | 0.65 | 0.464 | 2.05E-05 | 0 |
| FOXN3 | 1.56E-09 | 0.357 | 0.817 | 0.676 | 2.71E-05 | 0 |
| RGS10 | 1.74E-09 | 0.336 | 0.878 | 0.733 | 3.01E-05 | 0 |
| CD14 | 1.87E-09 | 0.31 | 0.714 | 0.479 | 3.24E-05 | 0 |
| ADAP2 | 2.32E-09 | 0.429 | 0.672 | 0.552 | 4.03E-05 | 0 |
| ABCC4 | 2.33E-09 | 0.393 | 0.399 | 0.218 | 4.05E-05 | 0 |
| KLF6 | 2.44E-09 | 0.702 | 0.958 | 0.885 | 4.23E-05 | 0 |
| MAFB | 3.33E-09 | 0.448 | 0.871 | 0.715 | 5.78E-05 | 0 |
| SGPP1 | 3.98E-09 | 0.342 | 0.566 | 0.358 | 6.92E-05 | 0 |
| LILRB4 | 4.11E-09 | 0.384 | 0.598 | 0.409 | 7.13E-05 | 0 |
| DDX5 | 4.13E-09 | 0.28 | 0.977 | 0.933 | 7.18E-05 | 0 |
| PLD4 | 6.14E-09 | 0.303 | 0.778 | 0.606 | 1.07E-04 | 0 |
| PIK3R1 | 9.89E-09 | 0.373 | 0.64 | 0.448 | 1.72E-04 | 0 |
| MAP3K8 | 1.11E-08 | 0.407 | 0.714 | 0.533 | 1.93E-04 | 0 |
| PXDC1 | 2.02E-08 | 0.286 | 0.408 | 0.215 | 3.51E-04 | 0 |
| FCHO2 | 2.17E-08 | 0.276 | 0.363 | 0.176 | 3.77E-04 | 0 |
| FAM105A | 2.20E-08 | 0.293 | 0.64 | 0.464 | 3.83E-04 | 0 |
| IL6ST | 2.54E-08 | 0.455 | 0.656 | 0.527 | 4.42E-04 | 0 |
| TGFBR1 | 2.82E-08 | 0.349 | 0.582 | 0.433 | 4.89E-04 | 0 |
| CCL3 | 2.82E-08 | 1.061 | 0.441 | 0.282 | 4.89E-04 | 0 |
| MVB12B | 3.21E-08 | 0.312 | 0.312 | 0.142 | 5.57E-04 | 0 |
| ITGB5 | 3.31E-08 | 0.329 | 0.392 | 0.218 | 5.74E-04 | 0 |
| TLR2 | 3.67E-08 | 0.332 | 0.688 | 0.521 | 6.36E-04 | 0 |
| PNRC1 | 4.12E-08 | 0.289 | 0.884 | 0.827 | 7.15E-04 | 0 |
| SIGLEC8 | 4.24E-08 | 0.286 | 0.273 | 0.112 | 7.36E-04 | 0 |
| TIMP2 | 4.43E-08 | 0.318 | 0.74 | 0.639 | 7.68E-04 | 0 |
| IRS2 | 4.60E-08 | 0.349 | 0.759 | 0.579 | 7.98E-04 | 0 |
| RUNX1 | 4.62E-08 | 0.334 | 0.45 | 0.27 | 8.03E-04 | 0 |

| | | | | | | |
|---|---|---|---|---|---|---|
| MYLIP | 4.97E-08 | 0.489 | 0.498 | 0.336 | 8.63E-04 | 0 |
| OLFML2B | 5.32E-08 | 0.374 | 0.28 | 0.121 | 9.23E-04 | 0 |
| LHFPL2 | 9.18E-08 | 0.291 | 0.412 | 0.227 | 1.59E-03 | 0 |
| EIF4A2 | 1.03E-07 | 0.332 | 0.842 | 0.77 | 1.79E-03 | 0 |
| HLA-DOA | 1.23E-07 | 0.302 | 0.73 | 0.6 | 2.14E-03 | 0 |
| LAPTM4A | 1.24E-07 | 0.299 | 0.707 | 0.585 | 2.15E-03 | 0 |
| DAGLB | 1.75E-07 | 0.362 | 0.498 | 0.342 | 3.03E-03 | 0 |
| TBXAS1 | 1.92E-07 | 0.283 | 0.733 | 0.624 | 3.33E-03 | 0 |
| AHSA2 | 2.87E-07 | 0.362 | 0.514 | 0.37 | 4.98E-03 | 0 |
| LPAR5 | 3.02E-07 | 0.296 | 0.37 | 0.203 | 5.25E-03 | 0 |
| C6orf62 | 3.10E-07 | 0.299 | 0.788 | 0.718 | 5.38E-03 | 0 |
| PIK3IP1 | 3.12E-07 | 0.313 | 0.331 | 0.173 | 5.42E-03 | 0 |
| EGR1 | 3.76E-07 | 0.585 | 0.791 | 0.761 | 6.52E-03 | 0 |
| FOXO3 | 4.34E-07 | 0.285 | 0.505 | 0.355 | 7.53E-03 | 0 |
| SOLH | 4.70E-07 | 0.362 | 0.608 | 0.482 | 8.16E-03 | 0 |
| KIAA1147 | 4.73E-07 | 0.278 | 0.28 | 0.133 | 8.20E-03 | 0 |
| CHD7 | 5.11E-07 | 0.329 | 0.338 | 0.179 | 8.86E-03 | 0 |
| WSB1 | 5.66E-07 | 0.289 | 0.891 | 0.824 | 9.83E-03 | 0 |
| P2RY13 | 5.67E-07 | 0.321 | 0.817 | 0.73 | 9.83E-03 | 0 |
| TLR7 | 6.31E-07 | 0.319 | 0.588 | 0.436 | 1.09E-02 | 0 |
| INPP5D | 6.95E-07 | 0.289 | 0.81 | 0.697 | 1.21E-02 | 0 |
| RNASE6 | 7.09E-07 | 0.305 | 0.83 | 0.709 | 1.23E-02 | 0 |
| LAIR1 | 7.49E-07 | 0.262 | 0.81 | 0.718 | 1.30E-02 | 0 |
| KIAA0247 | 7.68E-07 | 0.336 | 0.685 | 0.594 | 1.33E-02 | 0 |
| CTTNBP2NL | 8.47E-07 | 0.381 | 0.424 | 0.291 | 1.47E-02 | 0 |
| FMNL2 | 8.68E-07 | 0.306 | 0.37 | 0.209 | 1.51E-02 | 0 |
| SAP25 | 8.76E-07 | 0.303 | 0.344 | 0.197 | 1.52E-02 | 0 |
| RNF13 | 1.25E-06 | 0.274 | 0.646 | 0.512 | 2.17E-02 | 0 |
| CCL4 | 1.33E-06 | 0.897 | 0.64 | 0.512 | 2.30E-02 | 0 |
| ARRDC3 | 1.45E-06 | 0.44 | 0.588 | 0.491 | 2.51E-02 | 0 |
| GIMAP8 | 1.59E-06 | 0.262 | 0.547 | 0.388 | 2.76E-02 | 0 |
| HLA-DRB6 | 1.64E-06 | 0.342 | 0.971 | 0.882 | 2.85E-02 | 0 |
| FEZ2 | 1.68E-06 | 0.402 | 0.505 | 0.364 | 2.92E-02 | 0 |
| TFEC | 2.38E-06 | 0.267 | 0.486 | 0.324 | 4.13E-02 | 0 |
| PCM1 | 2.40E-06 | 0.28 | 0.605 | 0.479 | 4.16E-02 | 0 |
| IL8 | 2.52E-06 | 0.57 | 0.582 | 0.421 | 4.38E-02 | 0 |
| LRP1 | 2.79E-06 | 0.264 | 0.508 | 0.336 | 4.85E-02 | 0 |
| CD1C | 2.39E-52 | 1.491 | 0.808 | 0.219 | 4.14E-48 | 1 |
| CD1E | 3.53E-52 | 1.213 | 0.737 | 0.148 | 6.12E-48 | 1 |
| SLC38A1 | 4.74E-47 | 0.94 | 0.701 | 0.131 | 8.22E-43 | 1 |
| PARM1 | 1.25E-42 | 0.611 | 0.563 | 0.072 | 2.16E-38 | 1 |
| PLAC8 | 1.04E-39 | 1.035 | 0.856 | 0.333 | 1.80E-35 | 1 |
| PKIB | 1.93E-33 | 0.87 | 0.82 | 0.35 | 3.35E-29 | 1 |

| | | | | | | |
|---|---|---|---|---|---|---|
| FCER1A | 6.30E-32 | 0.921 | 0.976 | 0.601 | 1.09E-27 | 1 |
| PPA1 | 1.29E-29 | 0.809 | 0.886 | 0.479 | 2.24E-25 | 1 |
| KCNK17 | 1.48E-29 | 0.668 | 0.545 | 0.116 | 2.56E-25 | 1 |
| DENND1B | 1.97E-28 | 0.611 | 0.743 | 0.264 | 3.42E-24 | 1 |
| PAK1 | 6.82E-28 | 0.6 | 0.928 | 0.544 | 1.18E-23 | 1 |
| CD207 | 4.91E-26 | 1.39 | 0.796 | 0.468 | 8.52E-22 | 1 |
| GNAO1 | 5.11E-26 | 0.462 | 0.341 | 0.038 | 8.87E-22 | 1 |
| ADAM8 | 6.50E-26 | 0.531 | 0.641 | 0.196 | 1.13E-21 | 1 |
| PIK3R6 | 2.41E-24 | 0.59 | 0.581 | 0.177 | 4.18E-20 | 1 |
| STK17A | 5.29E-24 | 0.608 | 0.701 | 0.278 | 9.18E-20 | 1 |
| NAP1L1 | 7.31E-24 | 0.628 | 0.994 | 0.888 | 1.27E-19 | 1 |
| CDH17 | 1.73E-23 | 0.847 | 0.533 | 0.165 | 3.00E-19 | 1 |
| ASB2 | 4.01E-23 | 0.323 | 0.341 | 0.046 | 6.96E-19 | 1 |
| FLT3 | 1.03E-22 | 0.389 | 0.353 | 0.053 | 1.78E-18 | 1 |
| NAPSB | 2.77E-22 | 0.541 | 0.952 | 0.612 | 4.80E-18 | 1 |
| LSP1 | 3.31E-22 | 0.572 | 0.886 | 0.464 | 5.74E-18 | 1 |
| DAPK2 | 4.25E-22 | 0.29 | 0.281 | 0.027 | 7.38E-18 | 1 |
| RPS29 | 4.69E-22 | 0.378 | 1 | 0.992 | 8.15E-18 | 1 |
| RPS3A | 4.94E-22 | 0.352 | 1 | 0.998 | 8.58E-18 | 1 |
| CNN2 | 7.16E-22 | 0.525 | 0.946 | 0.618 | 1.24E-17 | 1 |
| SPATS2L | 9.23E-22 | 0.532 | 0.695 | 0.289 | 1.60E-17 | 1 |
| ST14 | 5.24E-21 | 0.409 | 0.737 | 0.281 | 9.10E-17 | 1 |
| BASP1 | 1.08E-20 | 0.548 | 0.952 | 0.595 | 1.87E-16 | 1 |
| AMICA1 | 1.74E-20 | 0.645 | 0.832 | 0.492 | 3.02E-16 | 1 |
| COTL1 | 5.13E-20 | 0.476 | 1 | 0.93 | 8.91E-16 | 1 |
| ADAM19 | 7.69E-20 | 0.329 | 0.365 | 0.07 | 1.33E-15 | 1 |
| RPL4 | 8.32E-20 | 0.373 | 0.994 | 0.968 | 1.44E-15 | 1 |
| PON2 | 3.61E-19 | 0.392 | 0.515 | 0.156 | 6.27E-15 | 1 |
| PRKAR2B | 6.08E-19 | 0.336 | 0.509 | 0.154 | 1.05E-14 | 1 |
| AFF3 | 8.76E-19 | 0.366 | 0.545 | 0.175 | 1.52E-14 | 1 |
| PABPC1 | 1.29E-18 | 0.292 | 1 | 0.998 | 2.24E-14 | 1 |
| CCR6 | 1.29E-18 | 0.473 | 0.527 | 0.175 | 2.24E-14 | 1 |
| RPS11 | 1.55E-18 | 0.329 | 1 | 1 | 2.69E-14 | 1 |
| MYO1D | 2.29E-18 | 0.284 | 0.329 | 0.061 | 3.97E-14 | 1 |
| RPL6 | 2.59E-18 | 0.329 | 0.994 | 0.958 | 4.49E-14 | 1 |
| MCOLN2 | 2.97E-18 | 0.478 | 0.599 | 0.236 | 5.15E-14 | 1 |
| EIF4G3 | 3.12E-18 | 0.387 | 0.659 | 0.266 | 5.42E-14 | 1 |
| RPL38 | 5.38E-18 | 0.341 | 1 | 0.981 | 9.34E-14 | 1 |
| AGPAT9 | 6.51E-18 | 0.481 | 0.611 | 0.243 | 1.13E-13 | 1 |
| CD101 | 7.25E-18 | 0.382 | 0.491 | 0.15 | 1.26E-13 | 1 |
| HIC1 | 8.30E-18 | 0.439 | 0.563 | 0.2 | 1.44E-13 | 1 |
| CLEC10A | 1.00E-17 | 0.65 | 0.695 | 0.331 | 1.74E-13 | 1 |
| ACTN1 | 1.43E-17 | 0.519 | 0.868 | 0.517 | 2.48E-13 | 1 |

| | | | | | | |
|---|---|---|---|---|---|---|
| ACTG1 | 1.85E-17 | 0.377 | 1 | 0.994 | 3.20E-13 | 1 |
| ATP2A3 | 2.01E-17 | 0.296 | 0.377 | 0.086 | 3.48E-13 | 1 |
| SEMA4C | 4.08E-17 | 0.335 | 0.659 | 0.249 | 7.09E-13 | 1 |
| APOBR | 5.20E-17 | 0.44 | 0.731 | 0.35 | 9.02E-13 | 1 |
| ARHGAP5 | 6.94E-17 | 0.371 | 0.671 | 0.278 | 1.21E-12 | 1 |
| TLR10 | 8.60E-17 | 0.49 | 0.796 | 0.42 | 1.49E-12 | 1 |
| FCGR2B | 9.26E-17 | 0.529 | 0.611 | 0.241 | 1.61E-12 | 1 |
| RPS21 | 9.28E-17 | 0.353 | 1 | 0.964 | 1.61E-12 | 1 |
| LCP1 | 1.44E-16 | 0.419 | 0.994 | 0.916 | 2.50E-12 | 1 |
| GDI2 | 1.60E-16 | 0.452 | 0.97 | 0.787 | 2.77E-12 | 1 |
| ITGB7 | 1.61E-16 | 0.385 | 0.497 | 0.165 | 2.80E-12 | 1 |
| RPS7 | 4.38E-16 | 0.32 | 1 | 0.981 | 7.61E-12 | 1 |
| SNHG5 | 7.13E-16 | 0.457 | 0.964 | 0.755 | 1.24E-11 | 1 |
| LTB | 7.95E-16 | 0.888 | 0.479 | 0.175 | 1.38E-11 | 1 |
| EEF1G | 1.38E-15 | 0.343 | 0.994 | 0.949 | 2.39E-11 | 1 |
| CD1D | 1.71E-15 | 0.383 | 0.683 | 0.293 | 2.97E-11 | 1 |
| RPS23 | 2.82E-15 | 0.292 | 1 | 0.996 | 4.90E-11 | 1 |
| NDRG2 | 4.37E-15 | 0.459 | 0.772 | 0.407 | 7.58E-11 | 1 |
| RPS14 | 4.48E-15 | 0.262 | 1 | 0.987 | 7.77E-11 | 1 |
| ACAP1 | 5.74E-15 | 0.281 | 0.413 | 0.12 | 9.96E-11 | 1 |
| SPINT2 | 7.48E-15 | 0.46 | 0.928 | 0.707 | 1.30E-10 | 1 |
| AIM1 | 7.60E-15 | 0.445 | 0.695 | 0.329 | 1.32E-10 | 1 |
| MLLT4 | 9.77E-15 | 0.314 | 0.491 | 0.173 | 1.70E-10 | 1 |
| HLA-DQA1 | 1.39E-14 | 0.356 | 1 | 0.966 | 2.42E-10 | 1 |
| CD1A | 1.46E-14 | 0.632 | 0.365 | 0.108 | 2.54E-10 | 1 |
| RPL23A | 1.90E-14 | 0.297 | 1 | 0.968 | 3.30E-10 | 1 |
| RPS24 | 2.09E-14 | 0.257 | 1 | 0.994 | 3.63E-10 | 1 |
| HMGN1 | 2.22E-14 | 0.388 | 0.97 | 0.698 | 3.86E-10 | 1 |
| IL16 | 4.25E-14 | 0.273 | 0.677 | 0.289 | 7.37E-10 | 1 |
| CD200R1 | 5.58E-14 | 0.256 | 0.341 | 0.093 | 9.68E-10 | 1 |
| NUP210 | 6.51E-14 | 0.263 | 0.443 | 0.148 | 1.13E-09 | 1 |
| GSN | 6.83E-14 | 0.439 | 0.94 | 0.738 | 1.18E-09 | 1 |
| CACNA2D3 | 7.00E-14 | 0.34 | 0.497 | 0.186 | 1.22E-09 | 1 |
| HLA-DQA2 | 9.96E-14 | 0.394 | 0.85 | 0.542 | 1.73E-09 | 1 |
| CYFIP2 | 1.06E-13 | 0.288 | 0.335 | 0.086 | 1.84E-09 | 1 |
| RPS18 | 1.13E-13 | 0.294 | 1 | 0.992 | 1.96E-09 | 1 |
| SUSD3 | 1.31E-13 | 0.305 | 0.605 | 0.259 | 2.27E-09 | 1 |
| Sep-06 | 2.17E-13 | 0.388 | 0.802 | 0.426 | 3.76E-09 | 1 |
| COMMD6 | 2.21E-13 | 0.343 | 0.898 | 0.635 | 3.83E-09 | 1 |
| RPL27A | 2.92E-13 | 0.257 | 1 | 0.987 | 5.07E-09 | 1 |
| NAV1 | 3.59E-13 | 0.334 | 0.695 | 0.329 | 6.24E-09 | 1 |
| ACTR3 | 5.39E-13 | 0.378 | 0.952 | 0.745 | 9.35E-09 | 1 |
| RPL28 | 5.47E-13 | 0.29 | 1 | 0.966 | 9.50E-09 | 1 |

| | | | | | | |
|---|---|---|---|---|---|---|
| RPL32 | 5.80E-13 | 0.262 | 1 | 0.994 | 1.01E-08 | 1 |
| ALCAM | 6.89E-13 | 0.357 | 0.784 | 0.42 | 1.20E-08 | 1 |
| MBOAT7 | 7.19E-13 | 0.386 | 0.647 | 0.325 | 1.25E-08 | 1 |
| RPS15A | 7.46E-13 | 0.277 | 1 | 0.983 | 1.30E-08 | 1 |
| AHR | 9.66E-13 | 0.426 | 0.766 | 0.443 | 1.68E-08 | 1 |
| LITAF | 1.03E-12 | 0.374 | 1 | 0.903 | 1.78E-08 | 1 |
| IRF4 | 1.29E-12 | 0.382 | 0.299 | 0.076 | 2.24E-08 | 1 |
| RPL26 | 1.32E-12 | 0.263 | 1 | 0.992 | 2.30E-08 | 1 |
| PALLD | 1.52E-12 | 0.303 | 0.635 | 0.3 | 2.63E-08 | 1 |
| HLA-DOB | 1.53E-12 | 0.352 | 0.419 | 0.15 | 2.66E-08 | 1 |
| RPL7 | 2.94E-12 | 0.266 | 0.994 | 0.973 | 5.11E-08 | 1 |
| HLA-DPB1 | 3.17E-12 | 0.295 | 1 | 0.987 | 5.50E-08 | 1 |
| ACTB | 3.94E-12 | 0.303 | 1 | 1 | 6.84E-08 | 1 |
| YWHAH | 4.68E-12 | 0.372 | 0.982 | 0.77 | 8.13E-08 | 1 |
| MAP4K1 | 4.77E-12 | 0.293 | 0.545 | 0.236 | 8.27E-08 | 1 |
| COPS3 | 6.35E-12 | 0.256 | 0.593 | 0.262 | 1.10E-07 | 1 |
| GCLC | 7.99E-12 | 0.269 | 0.713 | 0.371 | 1.39E-07 | 1 |
| GEM | 9.12E-12 | 0.414 | 0.353 | 0.116 | 1.58E-07 | 1 |
| RAB11FIP4 | 9.18E-12 | 0.294 | 0.569 | 0.264 | 1.59E-07 | 1 |
| RPL27 | 9.33E-12 | 0.282 | 1 | 0.964 | 1.62E-07 | 1 |
| RPL19 | 9.40E-12 | 0.254 | 1 | 0.983 | 1.63E-07 | 1 |
| NRARP | 1.28E-11 | 0.437 | 0.407 | 0.154 | 2.23E-07 | 1 |
| RPL35A | 1.40E-11 | 0.25 | 1 | 0.951 | 2.43E-07 | 1 |
| RPS4X | 1.48E-11 | 0.277 | 0.994 | 0.951 | 2.57E-07 | 1 |
| HINT1 | 1.95E-11 | 0.384 | 0.808 | 0.508 | 3.38E-07 | 1 |
| RPL37A | 2.60E-11 | 0.293 | 1 | 0.975 | 4.51E-07 | 1 |
| PHACTR2 | 2.80E-11 | 0.311 | 0.677 | 0.342 | 4.86E-07 | 1 |
| TMEM131 | 3.00E-11 | 0.28 | 0.617 | 0.289 | 5.20E-07 | 1 |
| RPL10A | 3.21E-11 | 0.265 | 1 | 0.973 | 5.57E-07 | 1 |
| MTR | 3.46E-11 | 0.276 | 0.611 | 0.295 | 6.00E-07 | 1 |
| PSTPIP2 | 3.62E-11 | 0.275 | 0.76 | 0.405 | 6.28E-07 | 1 |
| PLXNC1 | 3.97E-11 | 0.367 | 0.868 | 0.57 | 6.89E-07 | 1 |
| RAC2 | 4.97E-11 | 0.351 | 0.844 | 0.557 | 8.62E-07 | 1 |
| IL18 | 5.21E-11 | 0.334 | 0.91 | 0.635 | 9.05E-07 | 1 |
| SEL1L3 | 5.32E-11 | 0.264 | 0.623 | 0.304 | 9.24E-07 | 1 |
| ACOX1 | 7.40E-11 | 0.257 | 0.527 | 0.236 | 1.28E-06 | 1 |
| KCNMB1 | 8.66E-11 | 0.362 | 0.533 | 0.243 | 1.50E-06 | 1 |
| F11R | 9.37E-11 | 0.271 | 0.551 | 0.266 | 1.63E-06 | 1 |
| MAP2K1 | 1.02E-10 | 0.287 | 0.796 | 0.479 | 1.78E-06 | 1 |
| RPS5 | 1.21E-10 | 0.261 | 0.994 | 0.937 | 2.10E-06 | 1 |
| HPS5 | 1.22E-10 | 0.304 | 0.635 | 0.31 | 2.12E-06 | 1 |
| NET1 | 1.66E-10 | 0.319 | 0.497 | 0.219 | 2.88E-06 | 1 |
| H2AFY | 2.21E-10 | 0.298 | 0.976 | 0.8 | 3.83E-06 | 1 |

| | | | | | | |
|---|---|---|---|---|---|---|
| DUSP4 | 2.77E-10 | 0.463 | 0.275 | 0.082 | 4.81E-06 | 1 |
| PIP4K2A | 3.06E-10 | 0.298 | 0.784 | 0.473 | 5.31E-06 | 1 |
| REPIN1 | 3.34E-10 | 0.252 | 0.575 | 0.274 | 5.79E-06 | 1 |
| HMGCS1 | 3.42E-10 | 0.282 | 0.401 | 0.156 | 5.93E-06 | 1 |
| GOLGA8S | 3.71E-10 | 0.278 | 0.677 | 0.354 | 6.44E-06 | 1 |
| LPXN | 3.89E-10 | 0.328 | 0.832 | 0.542 | 6.75E-06 | 1 |
| PRCP | 4.14E-10 | 0.291 | 0.82 | 0.511 | 7.18E-06 | 1 |
| RPL31 | 4.16E-10 | 0.268 | 1 | 0.983 | 7.22E-06 | 1 |
| ATP2B4 | 5.28E-10 | 0.288 | 0.491 | 0.224 | 9.17E-06 | 1 |
| ARF6 | 5.89E-10 | 0.306 | 0.916 | 0.65 | 1.02E-05 | 1 |
| EEF1B2 | 6.08E-10 | 0.261 | 0.988 | 0.954 | 1.06E-05 | 1 |
| CD59 | 7.80E-10 | 0.325 | 0.611 | 0.335 | 1.35E-05 | 1 |
| DUSP5 | 9.25E-10 | 0.419 | 0.605 | 0.342 | 1.60E-05 | 1 |
| TMEM14C | 1.06E-09 | 0.347 | 0.713 | 0.445 | 1.85E-05 | 1 |
| SUB1 | 1.07E-09 | 0.276 | 0.88 | 0.618 | 1.86E-05 | 1 |
| TBC1D9 | 1.12E-09 | 0.321 | 0.772 | 0.473 | 1.95E-05 | 1 |
| PIGR | 1.23E-09 | 0.424 | 0.311 | 0.114 | 2.13E-05 | 1 |
| HLA-DQB2 | 1.38E-09 | 0.475 | 0.868 | 0.633 | 2.39E-05 | 1 |
| PCNA | 1.45E-09 | 0.409 | 0.473 | 0.215 | 2.51E-05 | 1 |
| GAPT | 1.61E-09 | 0.313 | 0.856 | 0.532 | 2.79E-05 | 1 |
| VASP | 1.78E-09 | 0.267 | 0.802 | 0.477 | 3.09E-05 | 1 |
| HLA-DQB1 | 1.84E-09 | 0.294 | 1 | 0.994 | 3.19E-05 | 1 |
| SLC25A5 | 1.94E-09 | 0.338 | 0.946 | 0.755 | 3.36E-05 | 1 |
| SSR1 | 2.30E-09 | 0.255 | 0.988 | 0.774 | 3.98E-05 | 1 |
| RUNX3 | 2.30E-09 | 0.38 | 0.82 | 0.544 | 3.99E-05 | 1 |
| SERTAD2 | 3.92E-09 | 0.288 | 0.629 | 0.344 | 6.81E-05 | 1 |
| LAMTOR1 | 4.20E-09 | 0.265 | 0.838 | 0.515 | 7.29E-05 | 1 |
| SNRPE | 4.96E-09 | 0.251 | 0.665 | 0.359 | 8.61E-05 | 1 |
| PSMB4 | 5.21E-09 | 0.289 | 0.838 | 0.532 | 9.04E-05 | 1 |
| TPM4 | 6.63E-09 | 0.29 | 0.916 | 0.633 | 1.15E-04 | 1 |
| CCND1 | 6.94E-09 | 0.295 | 0.515 | 0.253 | 1.20E-04 | 1 |
| EIF3L | 7.27E-09 | 0.267 | 0.97 | 0.795 | 1.26E-04 | 1 |
| L36A-HNRNP | 8.63E-09 | 0.313 | 0.964 | 0.806 | 1.50E-04 | 1 |
| STK17B | 9.36E-09 | 0.332 | 0.946 | 0.717 | 1.62E-04 | 1 |
| CKLF | 1.03E-08 | 0.286 | 0.82 | 0.513 | 1.79E-04 | 1 |
| ATP5L | 1.11E-08 | 0.272 | 0.958 | 0.715 | 1.93E-04 | 1 |
| DNTTIP2 | 1.13E-08 | 0.283 | 0.575 | 0.287 | 1.96E-04 | 1 |
| RPL13A | 1.48E-08 | 0.27 | 1 | 0.871 | 2.58E-04 | 1 |
| MBNL1 | 1.81E-08 | 0.259 | 0.922 | 0.741 | 3.14E-04 | 1 |
| VPS35 | 1.94E-08 | 0.27 | 0.904 | 0.643 | 3.36E-04 | 1 |
| UQCRB | 2.07E-08 | 0.267 | 0.862 | 0.536 | 3.59E-04 | 1 |
| BTF3 | 2.67E-08 | 0.263 | 0.976 | 0.789 | 4.63E-04 | 1 |
| CDK2AP2 | 2.71E-08 | 0.287 | 0.772 | 0.473 | 4.70E-04 | 1 |

| | | | | | | |
|---|---|---|---|---|---|---|
| HNRNPA1 | 3.37E-08 | 0.253 | 0.988 | 0.867 | 5.85E-04 | 1 |
| CIB1 | 3.85E-08 | 0.279 | 0.796 | 0.534 | 6.68E-04 | 1 |
| NKRF | 4.16E-08 | 0.297 | 0.473 | 0.23 | 7.23E-04 | 1 |
| HLA-DRB5 | 4.61E-08 | 0.294 | 0.802 | 0.563 | 8.00E-04 | 1 |
| FAM107B | 6.46E-08 | 0.296 | 0.569 | 0.316 | 1.12E-03 | 1 |
| PDLIM5 | 7.03E-08 | 0.27 | 0.647 | 0.386 | 1.22E-03 | 1 |
| PPHLN1 | 7.31E-08 | 0.273 | 0.581 | 0.319 | 1.27E-03 | 1 |
| IL2RG | 7.67E-08 | 0.257 | 0.754 | 0.456 | 1.33E-03 | 1 |
| PEBP1 | 8.58E-08 | 0.255 | 0.82 | 0.57 | 1.49E-03 | 1 |
| SNAP29 | 1.03E-07 | 0.264 | 0.641 | 0.39 | 1.78E-03 | 1 |
| HSPE1 | 1.35E-07 | 0.337 | 0.85 | 0.614 | 2.34E-03 | 1 |
| HIPK2 | 1.39E-07 | 0.253 | 0.731 | 0.468 | 2.41E-03 | 1 |
| NPM1 | 1.59E-07 | 0.256 | 0.982 | 0.8 | 2.76E-03 | 1 |
| HMGN2 | 1.76E-07 | 0.365 | 0.82 | 0.57 | 3.05E-03 | 1 |
| GLIPR1 | 5.01E-07 | 0.262 | 0.814 | 0.57 | 8.70E-03 | 1 |
| H2AFV | 5.51E-07 | 0.264 | 0.731 | 0.479 | 9.57E-03 | 1 |
| GZF1 | 1.05E-06 | 0.272 | 0.569 | 0.327 | 1.83E-02 | 1 |
| RPL17 | 1.11E-06 | 0.288 | 0.946 | 0.852 | 1.92E-02 | 1 |
| RPL36A | 2.39E-06 | 0.272 | 0.898 | 0.831 | 4.14E-02 | 1 |
| SLC38A2 | 2.67E-06 | 0.306 | 0.862 | 0.709 | 4.63E-02 | 1 |
| NFAT5 | 2.70E-06 | 0.255 | 0.533 | 0.308 | 4.68E-02 | 1 |
| S100A9 | 5.79E-65 | 2.567 | 0.626 | 0.038 | 1.01E-60 | 2 |
| S100A8 | 6.04E-57 | 2.409 | 0.589 | 0.048 | 1.05E-52 | 2 |
| FCN1 | 4.72E-55 | 2.175 | 0.773 | 0.199 | 8.19E-51 | 2 |
| VCAN | 3.24E-52 | 1.591 | 0.552 | 0.046 | 5.62E-48 | 2 |
| IFI30 | 2.21E-50 | 1.352 | 0.994 | 0.954 | 3.84E-46 | 2 |
| SH3BGRL3 | 4.83E-48 | 1.174 | 0.975 | 0.927 | 8.38E-44 | 2 |
| LYZ | 4.90E-41 | 1.78 | 0.957 | 0.805 | 8.51E-37 | 2 |
| SERPINA1 | 2.67E-40 | 1.16 | 0.926 | 0.676 | 4.63E-36 | 2 |
| S100A4 | 1.01E-31 | 1.368 | 0.871 | 0.573 | 1.75E-27 | 2 |
| PSAP | 1.33E-31 | 0.874 | 0.994 | 0.99 | 2.31E-27 | 2 |
| S100A10 | 7.55E-31 | 1.109 | 0.712 | 0.32 | 1.31E-26 | 2 |
| CD52 | 1.83E-29 | 1.317 | 0.528 | 0.144 | 3.17E-25 | 2 |
| TIMP1 | 5.47E-27 | 1.557 | 0.779 | 0.517 | 9.49E-23 | 2 |
| PRKCB | 3.18E-25 | 0.895 | 0.773 | 0.494 | 5.52E-21 | 2 |
| FLNA | 4.84E-25 | 0.948 | 0.638 | 0.295 | 8.40E-21 | 2 |
| PFN1 | 7.51E-25 | 0.543 | 0.994 | 0.996 | 1.30E-20 | 2 |
| HK3 | 1.43E-24 | 0.703 | 0.294 | 0.027 | 2.48E-20 | 2 |
| S100A6 | 4.09E-24 | 1.16 | 0.791 | 0.575 | 7.10E-20 | 2 |
| SLC11A1 | 1.45E-23 | 0.937 | 0.46 | 0.121 | 2.51E-19 | 2 |
| CFP | 2.59E-23 | 0.772 | 0.399 | 0.084 | 4.49E-19 | 2 |
| APOBEC3A | 1.55E-22 | 1.271 | 0.313 | 0.044 | 2.69E-18 | 2 |
| EMP3 | 4.46E-22 | 0.966 | 0.656 | 0.349 | 7.74E-18 | 2 |

| | | | | | | |
|---|---|---|---|---|---|---|
| LTA4H | 7.69E-22 | 1.008 | 0.706 | 0.412 | 1.33E-17 | 2 |
| TYMP | 1.11E-20 | 1.193 | 0.693 | 0.416 | 1.93E-16 | 2 |
| CTSS | 5.67E-20 | 0.744 | 0.951 | 0.964 | 9.85E-16 | 2 |
| FGR | 3.97E-19 | 0.889 | 0.834 | 0.653 | 6.88E-15 | 2 |
| HLA-B | 6.85E-19 | 0.483 | 0.988 | 1 | 1.19E-14 | 2 |
| CD36 | 1.89E-18 | 0.927 | 0.417 | 0.126 | 3.28E-14 | 2 |
| CSTA | 5.92E-18 | 0.79 | 0.546 | 0.238 | 1.03E-13 | 2 |
| FTH1 | 1.14E-17 | 0.671 | 1 | 1 | 1.99E-13 | 2 |
| MT2A | 2.17E-16 | 1.019 | 0.779 | 0.621 | 3.76E-12 | 2 |
| S100A11 | 2.85E-16 | 0.538 | 0.957 | 0.948 | 4.94E-12 | 2 |
| CD44 | 4.82E-16 | 0.785 | 0.699 | 0.504 | 8.37E-12 | 2 |
| TSPO | 6.76E-16 | 0.789 | 0.62 | 0.397 | 1.17E-11 | 2 |
| LY6E | 6.81E-16 | 0.721 | 0.46 | 0.178 | 1.18E-11 | 2 |
| THBS1 | 8.93E-16 | 1.333 | 0.307 | 0.073 | 1.55E-11 | 2 |
| MYL6 | 5.33E-15 | 0.489 | 0.939 | 0.893 | 9.26E-11 | 2 |
| NCF2 | 1.55E-14 | 0.813 | 0.675 | 0.469 | 2.69E-10 | 2 |
| ACTB | 3.62E-14 | 0.358 | 1 | 1 | 6.28E-10 | 2 |
| ANPEP | 6.78E-14 | 0.551 | 0.307 | 0.086 | 1.18E-09 | 2 |
| WARS | 2.01E-13 | 1.207 | 0.607 | 0.41 | 3.49E-09 | 2 |
| BIRC3 | 5.09E-13 | 0.969 | 0.503 | 0.253 | 8.83E-09 | 2 |
| ATP6V0C | 7.15E-13 | 0.513 | 0.89 | 0.877 | 1.24E-08 | 2 |
| CD300E | 8.11E-13 | 0.608 | 0.313 | 0.096 | 1.41E-08 | 2 |
| FPR1 | 9.57E-13 | 0.664 | 0.485 | 0.232 | 1.66E-08 | 2 |
| SOD2 | 1.04E-12 | 0.867 | 0.706 | 0.54 | 1.81E-08 | 2 |
| AGTRAP | 1.13E-12 | 0.502 | 0.294 | 0.084 | 1.95E-08 | 2 |
| CD97 | 1.21E-12 | 0.915 | 0.528 | 0.314 | 2.09E-08 | 2 |
| OAZ1 | 1.80E-12 | 0.376 | 0.963 | 0.973 | 3.13E-08 | 2 |
| TYROBP | 2.54E-12 | 0.272 | 0.988 | 0.998 | 4.41E-08 | 2 |
| LGALS3 | 3.67E-12 | 0.703 | 0.54 | 0.303 | 6.36E-08 | 2 |
| LILRB3 | 8.15E-12 | 0.695 | 0.552 | 0.339 | 1.41E-07 | 2 |
| ITGAL | 1.36E-11 | 0.656 | 0.528 | 0.305 | 2.36E-07 | 2 |
| SLC2A6 | 3.38E-11 | 0.547 | 0.344 | 0.132 | 5.86E-07 | 2 |
| SERF2 | 3.44E-11 | 0.385 | 0.939 | 0.941 | 5.97E-07 | 2 |
| DMXL2 | 3.50E-11 | 0.554 | 0.411 | 0.184 | 6.08E-07 | 2 |
| CEBPB | 4.11E-11 | 0.71 | 0.877 | 0.843 | 7.13E-07 | 2 |
| GAPDH | 4.90E-11 | 0.425 | 0.963 | 0.956 | 8.50E-07 | 2 |
| FKBP1A | 6.49E-11 | 0.53 | 0.828 | 0.774 | 1.13E-06 | 2 |
| FAM65B | 8.68E-11 | 0.807 | 0.503 | 0.316 | 1.51E-06 | 2 |
| LCP1 | 2.87E-10 | 0.387 | 0.957 | 0.929 | 4.98E-06 | 2 |
| BRI3 | 3.23E-10 | 0.564 | 0.767 | 0.741 | 5.61E-06 | 2 |
| SLC25A37 | 3.66E-10 | 0.613 | 0.387 | 0.178 | 6.36E-06 | 2 |
| ANXA2 | 4.27E-10 | 0.657 | 0.656 | 0.523 | 7.42E-06 | 2 |
| EHD1 | 4.39E-10 | 0.509 | 0.301 | 0.111 | 7.62E-06 | 2 |

| | | | | | | |
|---|---|---|---|---|---|---|
| LILRA6 | 6.18E-10 | 0.51 | 0.319 | 0.126 | 1.07E-05 | 2 |
| COTL1 | 6.37E-10 | 0.426 | 0.951 | 0.948 | 1.11E-05 | 2 |
| CFD | 6.82E-10 | 0.575 | 0.423 | 0.207 | 1.18E-05 | 2 |
| GLIPR2 | 9.53E-10 | 0.508 | 0.374 | 0.174 | 1.65E-05 | 2 |
| RHOA | 1.02E-09 | 0.366 | 0.902 | 0.923 | 1.77E-05 | 2 |
| FTL | 1.08E-09 | 0.369 | 0.982 | 1 | 1.87E-05 | 2 |
| HLA-A | 2.09E-09 | 0.295 | 0.969 | 0.998 | 3.63E-05 | 2 |
| SLC43A3 | 2.34E-09 | 0.508 | 0.399 | 0.199 | 4.07E-05 | 2 |
| IFITM2 | 4.63E-09 | 0.669 | 0.84 | 0.851 | 8.03E-05 | 2 |
| LILRB2 | 7.34E-09 | 0.628 | 0.724 | 0.642 | 1.27E-04 | 2 |
| VDR | 8.39E-09 | 0.532 | 0.252 | 0.09 | 1.46E-04 | 2 |
| SECTM1 | 9.84E-09 | 0.657 | 0.472 | 0.293 | 1.71E-04 | 2 |
| TMSB10 | 1.09E-08 | 0.321 | 0.988 | 0.992 | 1.89E-04 | 2 |
| SAMSN1 | 1.10E-08 | 0.552 | 0.534 | 0.339 | 1.91E-04 | 2 |
| TPM3 | 1.57E-08 | 0.371 | 0.926 | 0.879 | 2.73E-04 | 2 |
| SOCS3 | 1.86E-08 | 0.812 | 0.62 | 0.49 | 3.23E-04 | 2 |
| ZYX | 2.24E-08 | 0.438 | 0.877 | 0.862 | 3.88E-04 | 2 |
| AP2S1 | 2.39E-08 | 0.491 | 0.675 | 0.598 | 4.15E-04 | 2 |
| PSME2 | 2.54E-08 | 0.521 | 0.681 | 0.613 | 4.41E-04 | 2 |
| MYO1G | 3.40E-08 | 0.531 | 0.319 | 0.149 | 5.91E-04 | 2 |
| ALDOA | 5.07E-08 | 0.645 | 0.626 | 0.54 | 8.81E-04 | 2 |
| CD48 | 5.80E-08 | 0.551 | 0.681 | 0.59 | 1.01E-03 | 2 |
| ENO1 | 6.34E-08 | 0.422 | 0.828 | 0.872 | 1.10E-03 | 2 |
| C1orf162 | 6.83E-08 | 0.544 | 0.767 | 0.661 | 1.19E-03 | 2 |
| GRINA | 8.37E-08 | 0.564 | 0.656 | 0.59 | 1.45E-03 | 2 |
| PLAUR | 9.32E-08 | 0.48 | 0.73 | 0.59 | 1.62E-03 | 2 |
| APOL6 | 1.08E-07 | 0.515 | 0.706 | 0.615 | 1.88E-03 | 2 |
| CPPED1 | 1.62E-07 | 0.482 | 0.564 | 0.429 | 2.81E-03 | 2 |
| YWHAZ | 1.74E-07 | 0.382 | 0.877 | 0.866 | 3.02E-03 | 2 |
| LFNG | 2.10E-07 | 0.373 | 0.276 | 0.115 | 3.64E-03 | 2 |
| TNFRSF1B | 2.36E-07 | 0.494 | 0.804 | 0.77 | 4.09E-03 | 2 |
| CRIP1 | 2.74E-07 | 0.592 | 0.313 | 0.155 | 4.76E-03 | 2 |
| EFHD2 | 3.57E-07 | 0.461 | 0.755 | 0.736 | 6.20E-03 | 2 |
| SEMA4A | 3.58E-07 | 0.348 | 0.27 | 0.115 | 6.22E-03 | 2 |
| SELL | 4.18E-07 | 0.646 | 0.356 | 0.19 | 7.26E-03 | 2 |
| CYBB | 4.88E-07 | 0.386 | 0.865 | 0.872 | 8.47E-03 | 2 |
| Sep-09 | 9.69E-07 | 0.621 | 0.552 | 0.448 | 1.68E-02 | 2 |
| TNFAIP2 | 1.08E-06 | 0.583 | 0.638 | 0.542 | 1.88E-02 | 2 |
| COX4I1 | 2.31E-06 | 0.285 | 0.853 | 0.9 | 4.01E-02 | 2 |
| VASP | 2.65E-06 | 0.377 | 0.656 | 0.529 | 4.60E-02 | 2 |

**Supplementary Table 1.** Clinicopathological characterization of prostates and cell numbers included in this study

| Patient | Gleason Score | Initial PSA | pT | Age | Plate | # of tumorous cells before RNA quality check | # of tumor-adjacent cells before RNA quality check | # of tumorous cells after RNA quality check | # of tumor-adjacent cells after RNA quality check | # of cells sequenced per patient |
|---|---|---|---|---|---|---|---|---|---|---|
| 1 | 3+3 | 9.3 | pT2 | 63 | 1-2 | 527 | 210 | 527 | 247 | 774 |
| 2 | 3+4 | 7.6 | pT3 | 74 | 3 | 384 | 0 | 384 | 0 | 384 |
| 3 | 3+4 | 8.8 | pT2 | 50 | 4-5 | 384 | 384 | 0 | 384 | 384 |
| 4 | 4+3 | 8.3 | pT2 | 65 | 7-8 | 360 | 378 | 0 | 378 | 378 |
| | | Mean PSA | | Mean Age | | | | | | Total cells |
| Mean/Total | | 8,5 | | 63 | | | | 911 | 1009 | 1920 |
| Cells remaining after filtering and QC | | | | | | | | | | 751 |
| Cells remaining after NK cell removal | | | | | | | | 363 | 278` | **641** |

**Supplementary Table 3.** Significant differentially expressed canonical M1 and M2 associated genes

| | gene | p_val | avg_logFC | pct.1 | pct.2 | p_val_adj | cluster |
|---|---|---|---|---|---|---|---|
| M1 markers | BIRC3 | 5.09e-13 | 0.969 | 0.503 | 0.253 | 8.83e-09 | 2 |
| | SLC2A6 | 3.38e-11 | 0.547 | 0.344 | 0.132 | 5.86e-07 | 2 |
| | PSME2 | 2.54e-08 | 0.521 | 0.681 | 0.613 | 4.41e-04 | 2 |
| M2 markers | MAF | 2.08e-43 | 1.188 | 0.774 | 0.302 | 3.60e-39 | 0 |
| | MSR1 | 2.80e-17 | 0.545 | 0.698 | 0.425 | 4.86e-13 | 0 |
| | TGFBR2 | 2.48e-12 | 0.498 | 0.695 | 0.502 | 4.31e-08 | 0 |
| | SLC4A7 | 4.50e-12 | 0.461 | 0.666 | 0.458 | 7.81e-08 | 0 |
| | IGF1 | 6.04e-11 | 0.453 | 0.269 | 0.08 | 1.05e-06 | 0 |
| | MS4A4A | 1.51e-10 | 0.484 | 0.613 | 0.409 | 2.62e-06 | 0 |
| | CD36 | 1.08e-21 | 0.974 | 0.452 | 0.122 | 1.88e-17 | 2 |
| | LTA4H | 2.06e-21 | 0.984 | 0.71 | 0.411 | 3.58e-17 | 2 |

**Supplementary Table 4.** Summary of patient cohorts used for gene signature validation

| | Prospective GRID 5k | Mayo Clinic cohort | Johns Hopkins Medical Institutions cohort |
|---|---|---|---|
| | No. (%); Median (IQR) | No. (%); Median (IQR) | No. (%); Median (IQR) |
| Total | 5,239 (100%) | 780 | 355 |
| Age (years) (at RP or Bx) | 65.5 (60, 69.2) | 66(60,70) | 59(56,64) |
| PSA at diagnosis (ng/mL) | 6.5 (4.8, 9.7) | 9.4(6.1,18) | 8.6(5.7,13.3) |
| <10 ng/mL | 1886 (36%) | 410 (52.5%) | 209 (58.9%) |
| 10-20 ng/mL | 441 (8.4%) | 179 (22.9 %) | 111 (31.2%) |
| >20 ng/mL | 166 (3.1%) | 176 (22.5 %) | 34 (9.6%) |
| Gleason Grade group (pathology) | | | |
| Group 1 (GS 3+3) | 271 (5.1%) | 81 (10.4%) | 7 (2%) |
| Group 2 (GS 3+4) | 1769 (33.7%) | 390 (50%) | 150 (42.2%) |
| Group 3 (GS 4+3) | 1209 (23%) | NA | 66 (18.6%) |
| Group 4 (GS 8) | 396 (7.5%) | 107 (13.7%) | 35 (9.9%) |
| Group 5 (GS 9-10) | 554 (10.5%) | 202 (25.9%) | 97 (27.3%) |
| SM (positive) | 2099 (40%) | 401 (51.4%) | 102 (28.7%) |
| EPE (present) | 2092 (40%) | 372 (47.7%) | 238 (67%) |
| SVI (present) | 781 (15%) | 260 (33.3%) | 86 (24.2%) |
| LNI (positive) | 195 (3.7%) | 106 (13.6%) | 62 (17.5%) |
| Metastasis outcome | NA | 288 (36.9%) | 127 (35.8%) |
| High genomic risk (Decipher) | 1476 (28%) | 194 (24.8%) | 46 (12.9%) |
| Median follow-up (months) | 48 [36-54] | 156[106-201] | 108 [72-144] |
